# Structural insights into inhibitor mechanisms on immature HIV-1 Gag lattice revealed by high-resolution *in situ* single-particle cryo-EM

**DOI:** 10.1101/2024.10.09.617473

**Authors:** Chunxiang Wu, Megan E. Meuser, Juan S. Rey, Hamed Meshkin, Rachel Yang, Swapnil Chandrakant Devarkar, Christian Freniere, Jiong Shi, Christopher Aiken, Juan R. Perilla, Yong Xiong

**Author notes:** Correspondence (J.R.P.), (Y.X.). Equal contributions (Chunxiang Wu, Megan E. Meuser).

## Abstract

HIV-1 inhibitors, such as Bevirimat (BVM) and Lenacapavir (LEN), block the production and maturation of infectious virions. However, their mechanisms remain unclear due to the absence of high-resolution structures for BVM complexes and LEN’s structural data being limited to the mature capsid. Utilizing perforated virus-like particles (VLPs) produced from mammalian cells, we developed an approach to determine *in situ* cryo-electron microscopy (cryo-EM) structures of HIV-1 with inhibitors. This allowed for the first structural determination of the native immature HIV-1 particle with BVM and LEN bound inside the VLPs at high resolutions. Our findings offer a more accurate model of BVM engaging the Gag lattice and, importantly, demonstrate that LEN not only binds the mature capsid but also targets the immature lattice in a distinct manner. The binding of LEN induces a conformational change in the capsid protein (CA) region and alters the architecture of the Gag lattice, which may affect the maturation process. These insights expand our understanding of the inhibitory mechanisms of BVM and LEN on HIV-1 and provide valuable clues for the design of future inhibitors.

## Introduction

Production of an infectious virion is essential for the HIV-1 viral lifecycle, with the Gag polyprotein being a key player in virus production. The Gag polyprotein, composed of MA, CA, SP1, NC, SP2, and P6 domains, is the sole viral component needed for the assembly and release of an immature virus particle, which is accomplished by the oligomerization of Gag into a lattice structure at the plasma membrane^1^. Cryo-electron tomography (cryo-ET) data has revealed that the immature Gag lattice forms a spherical cage of interconnected hexamers where the CA protein plays a critical role in stabilizing the spherical hexameric lattice before the viral protease cleaves the Gag polyprotein into its individual subunits and maturation occurs^2–5^. CA is composed of two domains, the N-terminal domain (NTD) and the C-terminal domain (CTD). In the mature CA lattice, the NTD-NTD interactions help to stabilize neighboring protomers within a CA hexamer and CTD-CTD interactions stabilize inter-hexamer interactions. By contrast, the immature CA hexamer is stabilized by both CTD-CTD and NTD-NTD interactions and adopts a more extended conformation with a six-helix bundle at the base of the CA-SP1 domains, and the immature lattice is stabilized by NTD-NTD interactions that mediate the inter-hexamer contacts^3,6,7^. The immature HIV-1 Gag lattice is also known to interact with inositol hexakisphosphate (IP6) at the K158 and K227 rings within the central pore of a CA hexamer^7–12^. Recent studies have shown that the immature Gag lattice is critical for enriching IP6 into the HIV-1 virions for the proper maturation of the CA lattice^8^. It is important to note that IP6 is not required for the assembly of an immature Gag lattice, but the enriched IP6 level in virions is required for the proper assembly of the mature capsid. The proper formation of the interactions within the immature lattice is a crucial step for the overall success of the viral life cycle.

Due to the critical functions throughout the HIV-1 life cycle, the capsid has become a very attractive target for pharmacological inhibitors. An inhibitor of note is the first-generation maturation inhibitor, Bevirimat (BVM). BVM is known to stabilize the CA-SP1 six-helix bundle and prevent the final proteolytic cleavage of the Gag polyprotein, thereby preventing the proper maturation of HIV-1^13–16^. Previous studies using computational methods have indicated that BVM could bind the six-helix bundle in both BVM-up or BVM-down orientations, resulting in an ambiguous mechanism of binding for BVM^17^. Despite the advances with recent structural studies^18,19^, the exact binding pose and position of BVM have not been elucidated. Although BVM was highly effective at inhibiting HIV-1 initially, its activity was greatly reduced due to the frequent rise of resistance mutations^20–25^. However, since maturation inhibitors have great potential as candidates for HIV-1 treatment, the high-resolution structural information about how BVM binds to Gag plays an important role in the further development of this class of inhibitors.

Recently, a first-in-class highly potent inhibitor, Lenacapavir (LEN), also known as GS-6207, has received approvals for treating individuals with multi-drug resistant HIV-1 infections^26^. LEN is known to target the mature HIV-1 capsid by binding to the phenylalanine-glycine (FG)-binding pocket between an NTD and a CTD of two adjacent CAs in a hexamer and inhibit infection by altering the mechanical stability of the capsid shell and potentially interfering with CA-host protein interactions necessary for nuclear import, uncoating, and integration^27–29^. It has been reported that LEN also has a high binding affinity to Gag monomers, while treatment of LEN on HIV-1 infected cells caused a decreased amount of virus produced with mature morphologies and an increased amount of virus produced with aberrant capsid morphologies^28,29^. However, there is currently no known structure of LEN bound to immature CA, leaving a gap within the knowledge of the potential role of LEN during HIV-1 assembly and maturation.

To bridge the gap in understanding the binding modes and mechanisms of action imposed by inhibitors, we performed state-of-the-art *in situ* structural investigations into inhibitor interactions with the immature HIV-1 Gag lattice inside mammalian-produced virus-like particles (VLPs). We developed an effective approach to perforate the VLPs and determine *in situ* single-particle cryo-EM structures of the Gag lattice in complex with known inhibitors, including BVM and LEN. This enabled newfound insights into how these inhibitors work and demonstrated a successful approach for facilitating the design of future maturation inhibitors.

## Results

### High-resolution *in situ* structural analysis of the immature Gag_CA-SP1_ lattice assembly from intact and perforated VLPs

Recent works have shown that a single-particle cryo-EM approach can be used for high-resolution structural determination of large protein assemblies, like the HIV-1 CA lattice^30–32^. Here, we structurally characterized the immature HIV-1 CA-SP1 lattice (Gag_CA-SP1_) in a native environment by applying a single-particle cryo-EM approach directly on the Gag virus-like particles (VLPs) produced from mammalian cells (**Fig. 1a**). Additionally, to study the binding mechanisms of known HIV-1 inhibitors, we also resolved high-resolution structures of inhibitor-bound immature Gag_CA-SP1_ lattice assemblies (**Table 1**) from perforated VLPs, including BVM, LEN, or both of them bound to the VLPs (**Fig. 1**). These high-resolution *in situ* structures provide the most accurate depiction to date of BVM binding to immature HIV-1 particles. They also reveal a new mode of interaction, with LEN targeting the immature Gag assembly, in addition to its previously described binding to the mature capsid.

**Fig. 1.**
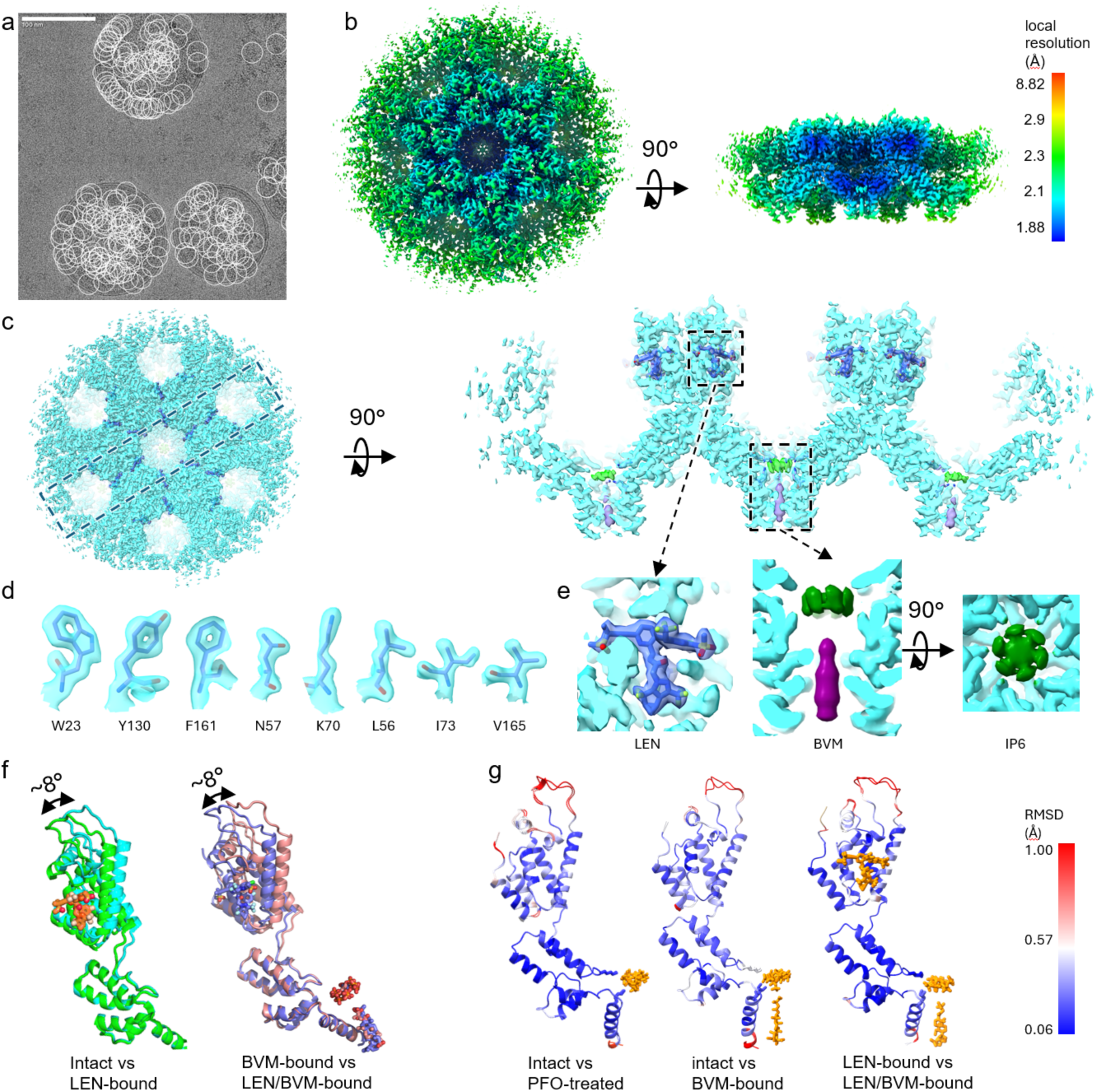
High-resolution *in situ* structural analysis of the immature Gag_CA-SP1_ lattice assembly from VLPs. a. An example micrograph of VLPs with picked single particles (white circles). The scale bar is 100 nm. b. Local resolution map of the immature Gag_CA-SP1_ lattice assembly with both LEN and BVM bound at 2.10 Å global resolution. The Gag_CA-SP1_ lattice assembly and ligands including LEN, BVM, and IP6 are well-resolved with high local resolution. c. The cryo-EM map for the immature Gag_CA-SP1_ lattice assembly (cyan) with both LEN and BVM bound from the top (left) and clipped side (right) views. The map regions for LEN, BVM, and IP6 are shown in royal blue, purple, and lime respectively. d. Examples of cryo-EM density of selected residues. e. Zoom-in view of the LEN, IP6, and BVM binding pockets with densities of LEN (royal blue), BVM (purple), and IP6 (green). f. LEN induces a conformational change that alters the relative orientation of the NTD to the CTD of CA by about 8°, compared to the forms without LEN bound in both intact/apo or BVM-bound VLPs. Left, alignment of immature Gag_CA-SP1_ protomers by superposition of the CTDs from VLPs without LEN (green, T8I VLP) and LEN-bound (cyan, NL-MA/NC VLP). Right, alignment of BVM-bound (slate, NL-MA/NC VLP) and LEN/BVM-bound (pink, T8I VLP) structures. g. Left, a comparison of models of intact T8I and PFO-treated NL-MA/NC VLPs shows that PFO treatment and the T8I mutation do not change the structure of Gag_CA-SP1_. Center and Right, comparisons of models of intact and BVM-bound (center) VLPs and models of LEN-bound and LEN/BVM-bound (right) VLPs show that BVM does not change the overall structure of Gag_CA-SP1_, with or without LEN bound.

**Table 1.** Cryo-EM data collection, refinement, and validation statistics.

|  | T8I Intact | T8I with PFO LEN/BVM-bound | T8I with PFO LEN-bound | NL-MA/NC with PFO | NL-MA/NC with PFO LEN-bound | NL-MA/NC with PFO BVM-bound |
| --- | --- | --- | --- | --- | --- | --- |
| EMDB | EMD-45975 | EMD-46593 | EMD-47240 | EMD-46631 | EMD-46594 | EMD-46595 |
| PDB | 9CWV | 9D6C | 9DWD | 9D88 | 9D6D | 9D6E |
| **Data collection and processing** |  |  |  |  |  |  |
| Magnification | 81,000 | 81,000 | 45,000 | 45,000 | 81,000 | 81,000 |
| Voltage (kV) | 300 | 300 | 200 | 200 | 300 | 300 |
| Electron exposure (e-/Å<sup>2</sup>) | 50 | 50 | 50 | 50 | 50 | 50 |
| Defocus range (μm) | 0.8 to 2.3 | 0.4 to 2.3 | 0.8 to 2.2 | 0.2 to 2.1 | 0.3 to 2.6 | 0.8 to 1.8 |
| Pixel size (Å) | 1.07 | 1.07 | 0.86 | 0.86 | 1.07 | 1.07 |
| Symmetry imposed | C6 | C6 | C6 | C6 | C6 | C6 |
| Initial particle images (no.) | 26,959,629 | 19,275,781 | 2,909,872 | 4,356,608 | 17,487,962 | 8,236,953 |
| Final particle images (no.) | 1,454,696 | 1,323,040 | 694,977 | 205,014 | 1,379,674 | 294,870 |
| Map resolution (Å) | 2.42 | 2.10 | 3.13 | 3.18 | 2.18 | 3.09 |
| FSC threshold | 0.143 | 0.143 | 0.143 | 0.143 | 0.143 | 0.143 |
| Map resolution range (Å) | 2.07-7.39 | 1.88-8.82 | 2.75-9.91 | 2.89-9.03 | 1.93-8.02 | 2.68-8.79 |
| **Refinement** |  |  |  |  |  |  |
| Initial model used (PDB code) | 7ASL | 9CWV | 9D6C | 9CWV | 9D6C | 9CWV |
| Model resolution (Å) | 2.5 | 2.1 | 3.3 | 3.3 | 2.3 | 3.2 |
| FSC threshold | 0.5 | 0.5 | 0.5 | 0.5 | 0.5 | 0.5 |
| Map sharpening B factor (Å<sup>2</sup>) | -14 | -24 | -42 | -61 | -33 | -71 |
| Model composition |  |  |  |  |  |  |
| Non-hydrogen atoms | 32976 | 34620 | 32832 | 31896 | 33792 | 31866 |
| Protein residues | 4158 | 4176 | 4122 | 4104 | 4128 | 4104 |
| Ligands | 7 | 32 | 19 | 1 | 25 | 2 |
| *B* factors (Å<sup>2</sup>) |  |  |  |  |  |  |
| Protein | 69 | 68 | 99 | 120 | 44 | 119 |
| Ligand | 149 | 89 | 117 | 168 | 61 | 137 |
| R.M.S. deviations |  |  |  |  |  |  |
| Bond lengths (Å) | 0.003 | 0.004 | 0.003 | 0.003 | 0.004 | 0.003 |
| Bond angles (°) | 0.6 | 0.7 | 0.6 | 0.6 | 0.7 | 0.5 |
| Validation |  |  |  |  |  |  |
| MolProbity score | 1.37 | 1.41 | 1.69 | 1.63 | 1.32 | 1.82 |
| Clashscore | 6.7 | 5.1 | 11.1 | 8.5 | 5.8 | 9.8 |
| Poor rotamers (%) | 0.9 | 1.5 | 1.5 | 1.6 | 0.6 | 2.5 |
| Ramachandran plot |  |  |  |  |  |  |
| Favored (%) | 99.0 | 99.0 | 98.6 | 98.6 | 98.4 | 98.9 |
| Allowed (%) | 1.0 | 1.0 | 1.4 | 1.4 | 1.6 | 1.1 |
| Disallowed (%) | 0 | 0 | 0 | 0 | 0 | 0 |

To produce the VLP samples, we initially utilized the Gag construct with a T8I mutation in the SP1 domain which is known to stabilize the immature lattice assembly by altering its structure ^7,21,33,34^. The VLPs were produced by transfecting mammalian HEK293T cells with a codon-optimized Gag expression vector and subsequently purified as previously described^7^. Additionally, we generated NL-MA/NC VLPs from a plasmid derived from the infectious molecular clone NL4-3 that contains point mutations at the protease cleavage sites between the MA, CA, SP1, and NC domains^35,36^. The NL-MA/NC VLPs were produced in Expi293F cells and purified for structural analysis. Using intact T8I VLPs, we were able to resolve the immature Gag_CA-SP1_ lattice structure to 2.42 Å resolution (**Table 1**). Our structure is consistent with previously published immature Gag_CA-SP1_ lattice structures from cryo-ET studies^3,6,7^, however, the high-resolution *in situ* structure that we obtained allowed us to reveal more atomic details at the assembly interfaces (described below)^7,37^.

To prepare the VLPs to facilitate the binding of inhibitors, we adopted a strategy to permeabilize the VLPs by a limited treatment with perfringolysin O (PFO)^38–40^, a cytolytic toxin that targets cholesterol-rich membranes, such as the lipid envelope of HIV-1, and forms ∼20-30 nm pores on the membrane bilayers^38^. The PFO treatment has been previously used on mature HIV-1 VLP samples for imaging analysis and cryo-ET structural studies of the mature HIV-1 capsid^41,42^, but has not yet been applied to immature VLPs. Our results show it is also a valuable tool for perforating immature VLPs while keeping the Gag lattice apparently unaltered. We used negative-stain EM to examine the level of permeabilization by PFO treatment. Before PFO treatment, only the membrane outlines of the VLPs could be stained and visualized without Gag lattice features in negative stain EM (**Fig. S1a)**. After successful pore formation by PFO treatment, the Gag lattice features became visible in negative stain (**Fig. S1b**). In cryo-EM micrographs, the Gag lattice remains mostly intact while some of the PFO-generated pores were visible (**Fig. S1c, S1d**). To confirm that limited PFO treatment does not alter the immature lattice structure, we solved the cryo-EM structure of PFO-treated NL-MA/NC VLPs, which showed a Gag_CA-SP1_ lattice structure identical to the one from intact T8I VLPs (**Table 1**, **Fig. 1g**). These results indicate that the limited PFO treatment can permeabilize the membrane of immature VLPs without disturbing the lattice structure, ideal for the study of Gag cofactors and inhibitors, especially those that are unable to diffuse through the viral membrane.

Using the perforated VLPs, we determined the cryo-EM structures of their complexes with BVM (3.09 Å resolution), LEN (2.18 Å resolution), or both BVM and LEN (2.10 Å resolution) (**Table 1**, **Fig. 1**). The high-resolution structures allowed for the confident modeling of the Gag_CA-SP1_ lattice structures and bound inhibitors (**Fig. 1c, 1d, 1e**). As previously reported^18,19^, the addition of BVM did not alter the Gag_CA-SP1_ structure or the immature lattice (**Fig. 1g**). However, LEN binding induced a conformational change in CA (**Fig. 1f**), accompanied by a change in the immature lattice architecture (described in detail below). The further addition of BVM to the LEN-bound VLP did not produce additional structural changes (**Fig. 1f, 1g**). These high-resolution structures demonstrate the effectiveness of the single-particle cryo-EM approach on PFO-treated VLPs for studying inhibitor structures of HIV particles.

### Detailed structural description of the immature Gag_CA-SP1_ lattice

Our immature Gag_CA-SP1_ lattice from the T8I intact VLPs is consistent with the previously published cryo-ET structures^7,43^, but the 2.42 Å resolution allowed us to analyze more interaction details of the immature lattice assembly. The immature lattice interfaces (**Fig. S2a**) include: (1) the intra-hexamer interface (1066 Å^2^ of buried surface area per protomer) between 2 adjacent protomers that are related in C6 symmetry; (2) the inter-hexamer interface (915 Å^2^ of buried surface area) between 2 C2-symmetrically related protomers; (3) the inter-hexamer interface (572 Å^2^ of buried surface area) between 2 C3-symmetrically related protomers.

Four major interfaces contribute to the interactions within the immature Gag_CA-SP1_ hexamer (**Fig. S2a**): (1) a top interface in the CA_NTD_, mediated by hydrogen bonds from R82 to Q50’ and T54’ and from E79 to N57’ (’ denotes the second chain); (2) a middle interface that is mediated by hydrogen bonds between S146 and the main chain carboxyl group of M144 to R173’; (3) a CA_CTD_ top interface that is mediated by hydrogen bonds between Q219’ and the main chain carboxyl groups of Q155 and N193; (4) a bottom interface that is mediated by hydrophobic interactions among V230, A228, L231 of the CA_CTD_ and M4, I8 of SP1.

For the inter-hexamer interfaces, our model depicts the details of two major interface areas in between two protomers with C3 symmetry (**Fig. S2a**): (1) a hydrogen bond from R18 to E76’ (’ marks residues from a different protomer), coupled with a nearby hydrophobic patch between residues T19, V26, K30, L43, M39 and residues L136’, I135’, I129’; (2) a hydrogen bond from E29 to R143’. Additionally, in between two interacting protomers with C2 symmetry, there are also two major interface areas (**Fig. S2a**): (1) R18 forms a hydrogen bond to the mainchain carboxyl group of N57’ and a salt bridge to T58’; N21 forms a hydrogen bond to the mainchain carboxyl group of L20’; (2) a hydrophobic patch mediated by A177, V181, W184, and M185. Given the C2 symmetry between the two chains, all the interface areas are symmetrically related as well.

It is interesting to note that in the immature Gag_CA-SP1_ lattice, R18, which mediates the IP6 binding pocket at the central pore of the mature CA hexamer, is at the critical interface between three adjacent immature Gag_CA-SP1_ protomers (**Fig. S2a**). The R18 residue forms hydrogen bonds with E76 from the C3-related chain and N57 from the C2-related chain (**Fig. S2a**). We also observed well-defined density for inositol hexakisphosphate (IP6), which is known to tightly bind to the K158/K227 rings of immature Gag_CA-SP1_ hexamers and help enrich IP6 into the virions^7,9,11,41,44^. With our improved resolution, we now can resolve the IP6 density to greater detail, including densities for a more precise conformation of the bound IP6 and the side chains of the lysine residues that coordinate the IP6 molecule (**Fig. S3**). Based on the model, the top K158 ring mainly contributes to the interaction with the 5 equatorial phosphates of the IP6, while the K227 ring mainly mediates the interaction with the axial phosphate. Nevertheless, since the asymmetric IP6 is not compatible with the C6 symmetry of the lattice, the final cryo-EM map for the IP6 is an average of the 6-fold symmetry (**Fig. S3**).

### Defining a new mode of LEN binding to the immature lattice

LEN is a recently FDA-approved CA inhibitor whose mechanism of action with the mature CA has been thoroughly characterized^27–29,45^. However, very few studies investigating the interaction of LEN with the immature Gag lattice have been performed. It is known that LEN can bind to the Gag monomer and treatment of virus-producing cells with LEN leads to virions with aberrant CA morphologies, but the structure of LEN bound to the immature Gag assemblies has yet to be elucidated^28,29^. We utilized the PFO-treated VLPs with either LEN or both BVM and LEN bound, which resulted in a 2.18 Å resolution and a 2.10 Å resolution cryo-EM maps, respectively (**Table 1**), with virtually identical Gag_CA-SP1_ conformations (RMSD 0.4 Å) (**Fig 1g**).

LEN binds the immature Gag_CA-SP1_ lattice in a novel binding pocket consisting of three different CA_NTD_ protomers, two from one hexamer (denoted chain #1 and #2) and the other from a neighboring hexamer (denoted chain #3) (**Fig. 1c, 3e, 2b, S2b**). This is a unique binding manner compared to that of LEN binding the mature CA lattice, which is composed of the NTD of one CA protomer (chain #1) and mostly the CTD of the adjacent CA protomer from the same hexamer (**Fig. 2a**). The unique binding environment leads to a conformational change in LEN when bound to the immature versus the mature CA lattices, although some similarities in the drug-binding pose are retained (**Fig. 2c**). The main binding pocket on chain #1 and the associated interactions are largely conserved. This interface includes helix 3 (residues P49-N57), the 58-TVGG-61 loop, and helix 4 (residues Q63-L83). The LEN-contacting residues, such as Thr54, Leu56, Asn57, Met66, Gln67, Lys70, Ile73, Tyr130, and Thr107 on this interface exhibit similar conformations and consistently maintain LEN in a similar pose in the mature and immature structures (**Fig. 2d**).

**Fig. 2.**
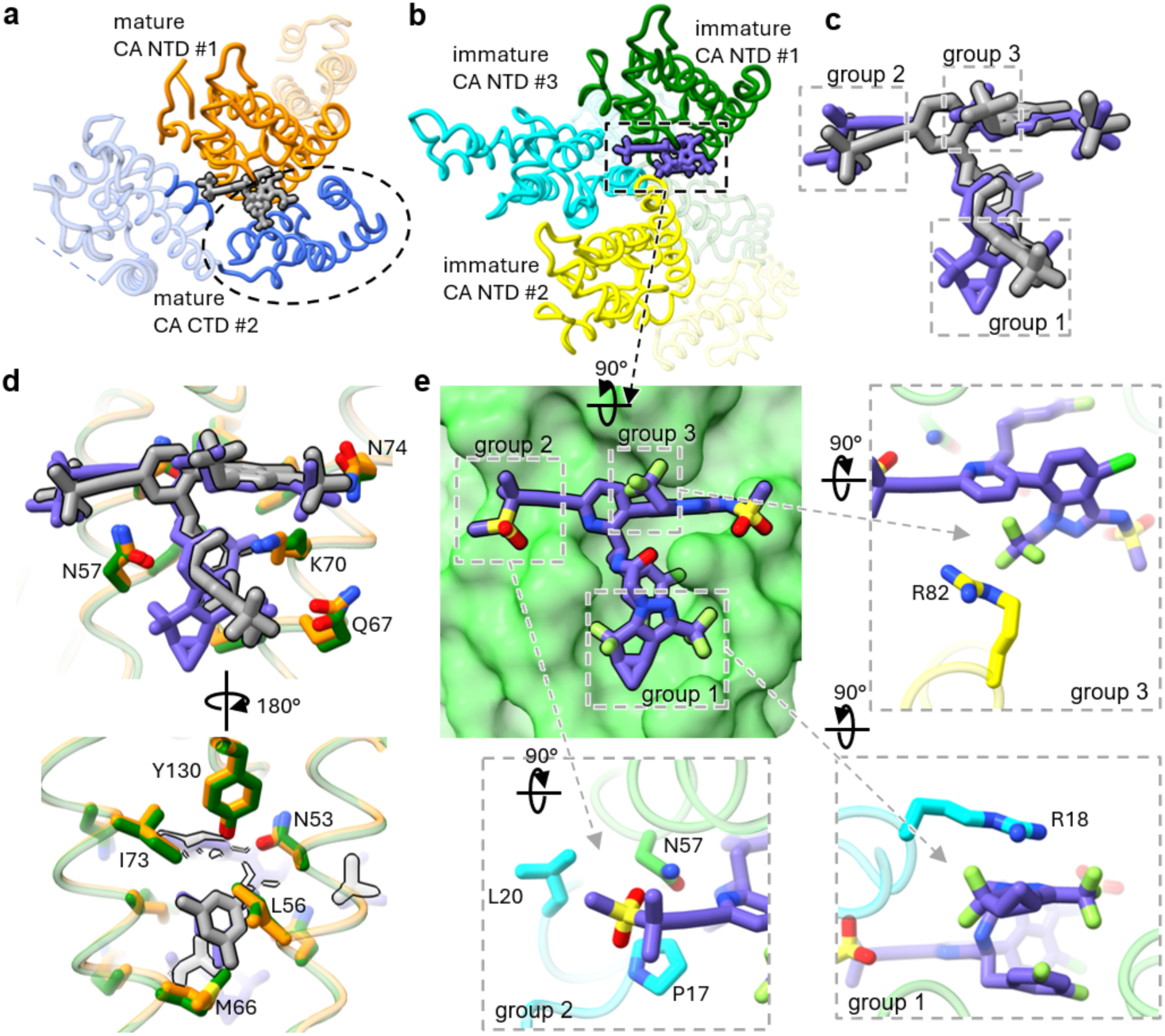
Comparison of LEN binding pocket in the immature and the mature CA lattices. a. LEN binding site in the mature capsid (PDB 6V2F). The binding site is mainly composed of the NTD of the major binder CA chain (orange) and the CTD of another CA chain (blue) within the same hexamer. LEN is shown in grey color. b. LEN binding site in the immature Gag_CA-SP1_ lattice. The binding site is mediated by NTDs only from three Gag_CA-SP1_ chains, including a major binder (green, oriented the same way as the major binder in a), a second chain within the same hexamer (yellow), and a third chain from an adjacent hexamer (cyan). LEN is shown in purple color. c. Alignment of the LEN conformations in the mature capsid (grey) and immature Gag_CA-SP1_ (purple) lattices. The regions engaged in unique interactions are boxed. d. Comparison of the LEN binding pocket at the major binding interface in the mature and the immature lattices. The mature CA is shown in orange, and the immature Gag_CA-SP1_ is shown in green. The residues with conserved interactions are shown as sticks. e. Unique interfaces at the LEN binding site in the immature Gag_CA-SP1_ lattice. The main binding interface is highlighted in green (top left), with additional unique interactions displayed in the insets. The methylsulfonylbutyl group of LEN interacts with L20 and P17 (cyan) from a Gag_CA-SP1_ chain of an adjacent immature hexamer (bottom left). The diazatricyclo-nonane group interacts with R18 (cyan) from the same adjacent immature hexamer (bottom right). Additionally, the trifluoroethyl group interacts with R82 (yellow) from the Gag_CA-SP1_ chain within the same hexamer as the major binder chain (top right).

**Fig. 3.**
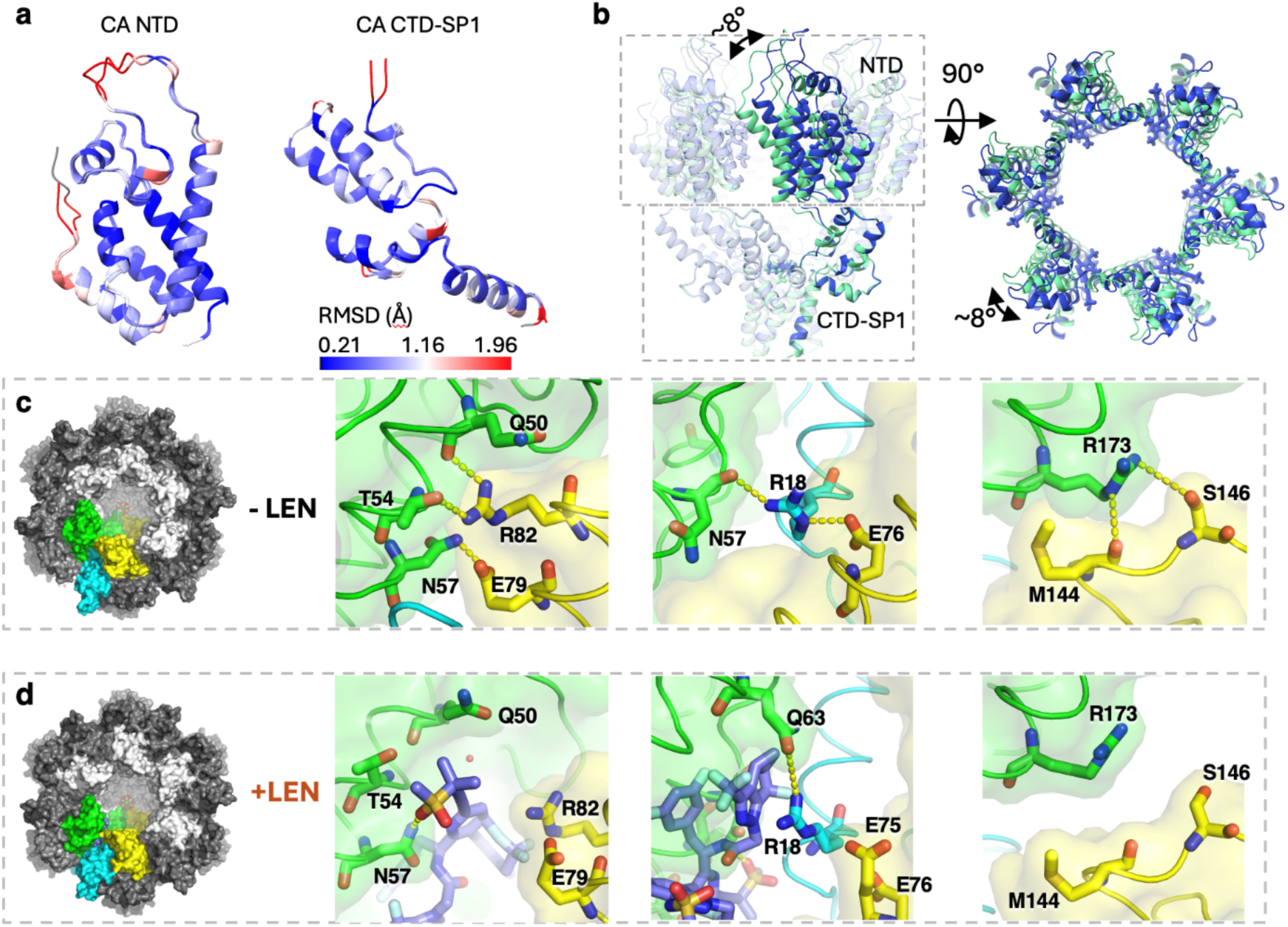
LEN binding induces conformational reorientation of the CA_NTD_. a. Alignment of the immature CA_NTD_ (left) and CA_CTD_-SP1 (right) from VLPs with or without LEN, which shows that the overall folding of the individual CA_NTD_ or CA_CTD_-SP1 is not changed upon LEN binding. The models are colored by the RMSD between the two Gag_CA-SP1_ structures. b. Illustration of the conformational change of immature Gag_CA-SP1_ hexamer before (lime green) and after (blue) LEN binding. The models of Gag_CA-SP1_ hexamers are aligned and displayed from the side view (left) and the CA_NTD_ is displayed from the top view (right). LEN induces an ∼8° rotation of the CA_NTD_ relative to the CA_CTD_. This conformational change rearranges the CA_NTD_ organization in the lattice assembly. c. Key interfaces that mediate the CA NTD-NTD interaction in the native Gag_CA-SP1_ lattice without LEN bound. Two CA-SP1 protomers in the same immature Gag_CA-SP1_ hexamer are colored in green and yellow respectively, while a third CA-SP1 protomer from an adjacent hexamer is colored in cyan. d. New interfaces that are induced by LEN binding in between the CA_NTD_ protomers. Three CA-SP1 protomers are colored the same as in 3c. The LEN bound at the CA NTD-NTD interface is colored in slate. At the corresponding interface compared to the native Gag_CA-SP1_ lattice, LEN reorganizes the hydrogen bond network and alters the orientation of the CA_NTD_.

By contrast, the additional, distinct CA_NTD_ interactions in the immature lattice (chains #2 and #3) cause conformational changes in LEN (**Fig 3e**). The largest conformational difference is a ∼90-degree rotation of the diazatricyclonona group of LEN when compared to the structure of LEN bound to the mature capsid (**Fig. 2a, 2b, 2c**). This change is likely due to the stacking interaction between the diazatricyclonona group of LEN and residue R18 of chain #3 from a neighboring CA hexamer (**Fig. 3e**). Notably, since R18 plays an essential role in coordinating IP6 in the mature capsid, the interaction between LEN and R18 may have further implications for capsid maturation. Furthermore, residues P17 and L20 on chain #3 create a binding pocket and alter the conformation of the methylsulfonylbut group of LEN. In addition, residue R82 on chain #2 of the same hexamer as chain #1 stacks onto the trifluoroethyl group of LEN, leading to a modest shift of its position. It is also noteworthy that some of the LEN-contacting residues previously observed in the mature CA lattice, such as S41 and Q179^27,31^, are no longer involved in the binding of LEN to the immature CA lattice. The unique binding of LEN across three CA_NTD_ also suggests that it modulates the NTD-NTD interactions between CA subunits both within and between immature hexamers.

### LEN binding induces changes in the architecture of the Gag_CA-SP1_ lattice

The binding of LEN induces a conformational change to the Gag_CA-SP1_ lattice structure, consistent with its multi-protomer binding site. This aligns with the observation of increased aberrant capsid morphologies in virions produced in the presence of LEN^29^. Alignment of the Gag_CA-SP1_ protomers with or without LEN by either CA_NTD_ or CA_CTD_ individually showed that no major changes occurred to the domain structures (**Fig. 3a**). However, an ∼8° reorientation of the CA_NTD_ relative to the CA_CTD_ takes place upon LEN binding (**Fig. 1f, 3b**), which induces an overall conformational change of the CA_NTD_ layer in the immature Gag_CA-SP1_ lattice (**Fig. 3b**) (**Supplemental Movies 1, 2**). By contrast, the interfaces in the CA_CTD_ remain unchanged (**Fig. S2b**), including the hydrophobic interfaces in the CA-SP1 six-helix bundle, regardless of whether additional binding with BVM occurs or not.

The change in the Gag_CA-SP1_ lattice is driven by the repositioning of residues to interact directly with LEN. When compared to the apo structures, the engagement of residues with LEN (**Fig. 2e**) results in the loss of certain interactions between protomers. Within an immature CA hexamer, hydrogen bonds are no longer formed between E79/R82 and Q50’/T54’/N57’ upon LEN binding (**Fig. 3c, 3d left**). In addition, due to the reorientation of the CA_NTD_, the hydrogen bonds between M144/S146 and R173’ are also lost (**Fig. 3c, 3d right**). Notably, residue R18 mediates the interactions at the juncture of three CA_NTD_ protomers and participates in the interaction with LEN, highlighting its key role in LEN binding and lattice dynamics. This residue is engaged in a hydrogen bond across the 2-fold interface with a neighboring E75’ in the LEN-bound structure instead of E76’ in the apo lattice (**Fig. 3c, 3d middle**). Similarly, R18 forms a new hydrogen bond across the 3-fold interface with Q63’ (LEN-bound) instead of N57’ (apo) (**Fig. 2e, 3c, 3d middle**).

### Defining the BVM binding position on the immature Gag_CA-SP1_ lattice

The maturation inhibitor BVM has been shown to bind at the CA-SP1 six-helix bundle of the Gag lattice by previous studies using microcrystal electron diffraction (MicroED) and magic angle spinning NMR (MAS-NMR)^18,19^. Both studies utilized *in vitro* assembled protein complexes with Gag truncations containing CA_CTD_-SP1. Furthermore, computational studies have proposed two potential BVM orientations based on the positioning of the dimethyl-succinate moiety^17,18^: “up-orientation” where the moiety points toward the CA_CTD_ and a “down-orientation” where the moiety points toward the SP1 domain. However, the molecular interactions underlying BVM binding remain controversial, as these studies suggest different binding poses of BVM. This was largely due to the difficulty of obtaining BVM-bound native Gag lattices for high-resolution structural work and the symmetry mismatch of the asymmetrical BVM molecule against the 6-fold symmetric immature Gag lattice. Due to these challenges, the binding pose of BVM has not yet been conclusively defined.

In this study, we present a more accurate BVM binding pose, determined through high-resolution in situ cryo-EM structures of the full-length Gag lattice within VLPs, complemented by full-atom molecular dynamics (MD) simulations. Utilizing PFO-treated VLPs, we obtained two BVM-bound immature Gag_CA-SP1_ structures, which include: (1) a 2.10 Å resolution structure with both LEN and BVM bound to T8I VLPs; (2) a 3.09 Å resolution BVM-bound structure from NL-MA/NC VLPs (**Table 1**). A comparison of the two structures reveals that BVM binding in both VLPs is consistent (**Fig. 4a, b**).

**Fig. 4.**
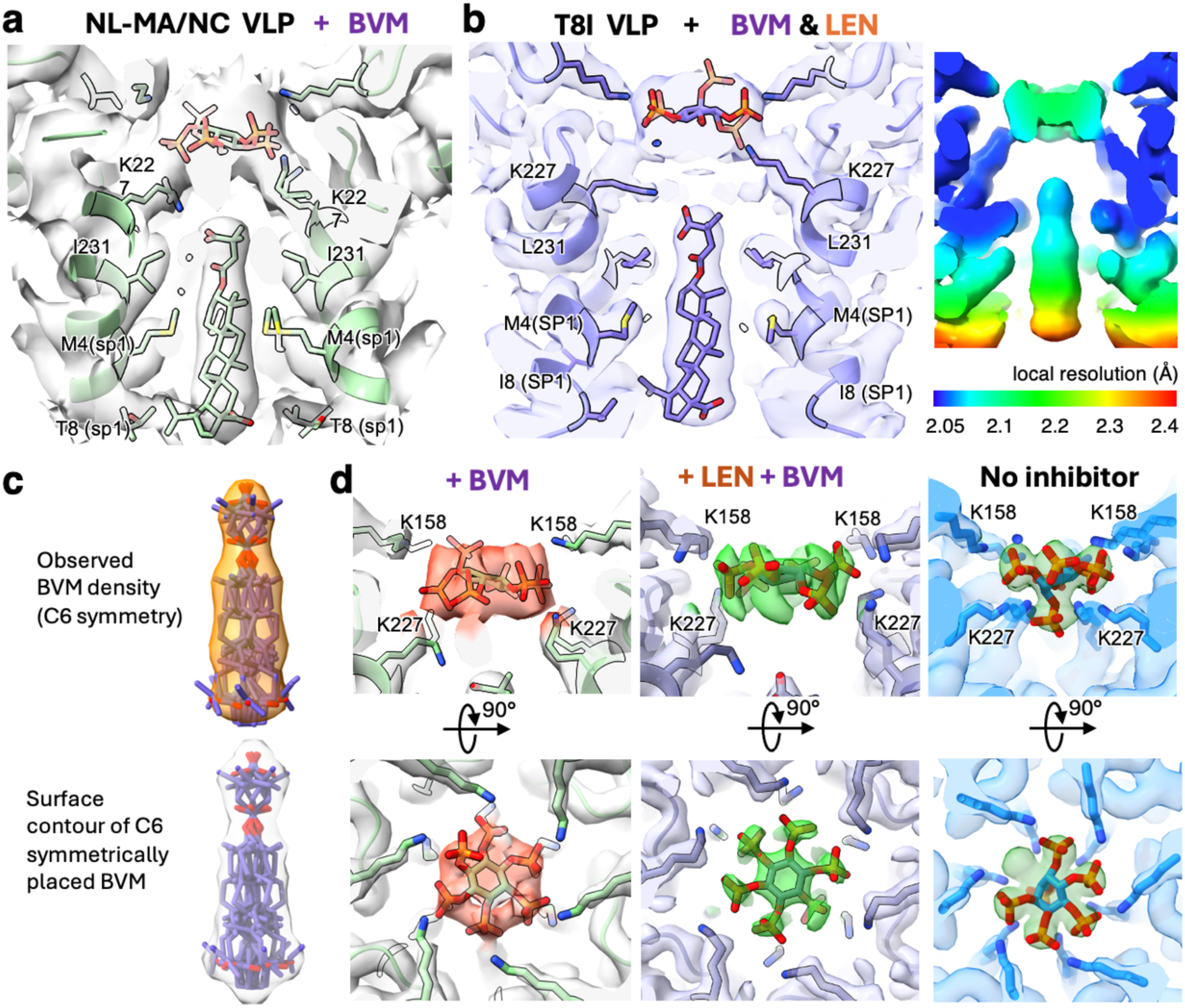
Binding poses of BVM and IP6 in the immature Gag_CA-SP1_ lattice. a. BVM binding in the immature Gag_CA-SP1_ lattice from PFO-treated NL-MA/NC VLPs. The cryo-EM density is shown in white and the atomic model is shown in green. IP6, BVM, and residues that form the binding pocket for BVM (K227, I231 of CA and M4, T8 of SP1) are shown in sticks. b. BVM binding in the immature Gag_CA-SP1_ lattice from PFO-treated T8I VLPs treated with both BVM and LEN. Left: The cryo-EM density and the atomic model are shown in slate color. IP6, BVM, and residues that form the binding pocket for BVM are shown in sticks. Right: Local resolution of the cryo-EM map. c. Fit of BVM models in the cryo-EM density. Top: The observed cryo-EM map is fitted with six copies of BVM, arranged with C6 symmetry. Bottom: For comparison, a computationally generated surface contour is shown, based on the six-copy BVM model with C6 symmetry. The surface was generated using ChimeraX^46^ “molmap” utility at a resolution of 3 Å. d. Change of the IP6 pose upon BVM binding. Side (upper panels) and top (lower panels) views of the IP6 density from PFO-perforated VLPs treated with BVM (left), PFO-perforated VLPs treated with BVM and LEN (middle), or intact VLPs without any inhibitor treatment or PFO perforation (right). The CA-SP1 residues interacting with IP6 are labeled.

Our high-resolution *in situ* cryo-EM study revealed a well-defined BVM density at the binding pocket with a local resolution range of 2.05 Å to 2.4 Å (**Fig. 4b**). Despite the high resolution, the density for BVM is an average of 6 symmetrically placed molecules since the Gag_CA-SP1_ lattice is six-fold symmetric (C6 symmetry) while the BVM bound at the symmetry center is not. To better interpret the averaged density, we modeled six symmetrically related BVM molecules inside the cryo-EM density (**Fig. 4c, left**), which was then compared with a surface contour generated from the symmetrically placed BVM molecules. This led to accurate positioning of the BVM model consistent with the “up-orientation,” where the dimethyl-succinate group is oriented up towards the CA_CTD_. However, the location of the bound BVM is more distant from the IP6 binding site and closer to the SP1 region than previous structural studies proposed^18,19^. Compared to the earlier studies, the current position of the succinyl moiety shows distance differences of approximately 5.4 Å relative to the microED structure and 9.7 Å relative to the NMR structure (**Fig. S6b, S6e**)^18,19^. The succinyl moiety is in the proximity of the K227 ring, which appears to adopt alternative conformations to interact with either IP6 or the BVM moiety (described in detail below). (**Fig. 4b**). In this binding mode, the betulinic acid moiety of BVM is positioned to interact with residues ranging from I231 of CACTD to T8 of SP1 in NL-MA/NC VLPs (**Fig. 4a**) or L231 to I8 in T8I VLPs (**Fig. 4b**).

### BVM potentially alters the conformation of bound IP6

Another distinct difference observed in the *in situ* structures is the altered shape of the density for IP6 upon BVM binding. IP6 is a known host-cell cofactor that promotes the assembly and maturation of CA lattice^8–10,12,44^. In the immature lattice, the binding of IP6 is mediated by the K158 and K227 rings within the CA-SP1 hexamer central pore, and analysis of our structures reveals that the IP6 molecule remains present in the K158/K227 rings regardless of inhibitor treatments. However, the pose of the bound IP6 molecule is altered when BVM is present (**Fig 4e**). In the apo structure, the IP6 molecule adopts a conformation where one of the phosphates in the axial position interacts with the K227 residues only and is positioned in the pore center of the six-helix bundle towards the SP1 domain, while the other equatorial phosphates radially interact with both K158 and K227 residues (**Fig. 4e** right). Upon BVM binding, the density for the axial phosphate of IP6 is no longer in the pore center. Instead, all phosphate moieties are positioned between equatorial and axial orientations toward the periphery, exhibiting increased flexibility to interact with both the K158 and K227 rings. (**Fig. 4a, 4d, 4e** left and right). Interestingly, the density for residue K227 suggests potential alternative side chain conformations (**Fig. 4a**), where K227 can potentially either interact with the succinyl moiety of BVM or engage the phosphate of IP6. Previous NMR study^19^ also showed that IP6 has a weakened interaction with K227 when BVM is bound. However, instead of IP6 moving away from the K227 ring and solely interacting with K158 residues as previously reported, we observed that IP6 remains positioned between the K158 and K227 rings at the canonical binding site upon BVM binding. The succinyl moiety of BVM may compete with the phosphate groups of IP6 for interaction with K227, resulting in an altered binding pose of IP6.

The binding of BVM does not alter the structure of Gag_CA-SP1_. Alignment of the Gag_CA-SP1_ models from the apo immature Gag lattice and the BVM-bound Gag lattice, showed an RMSD of ∼0.6 Å (RMSD ∼0.4 Å if excluding the flexible CypA-binding loop region spanning CA residues 88-94) **(Fig. 1g, middle)**. Similarly, the addition of BVM did not affect the structure of the LEN-bound Gag_CA-SP1_. A structural comparison between the Gag_CA-SP1_ model in complex with LEN and the model bound to both BVM and LEN showed an RMSD of ∼0.4 Å **(Fig. 1g, right**), demonstrating that BVM does not induce any additional conformational change in the LEN-bound Gag_CA-SP1_. Together, these results show that BVM binding does not affect the conformation of Gag_CA-SP1_ and that this effect is independent of the structural impact caused by LEN binding. This structure preservation is consistent with the previous microED structural study^18^ (**Fig. S6a,6b**) but contrasts slightly with previous NMR findings, which suggested that BVM binding induces a tightening of the SP1 six-helix bundle^19^ (**Fig. S6c**). This difference may be attributed to variations in Gag constructs and experimental conditions, specifically the use of *in situ* VLPs in this study compared to the in vitro assembled CA_CTD_-SP1 particles used previously, which may also contribute to the difference in the BVM binding location (**Fig. S6b, S6e**).

### *In silico* characterization of BVM binding

To better define the BVM binding mode that was averaged by the symmetric lattice of Gag_CA-SP1_, we conducted a *de novo in silico* characterization using molecular dynamics simulations (MD) of the CA_CTD_-SP1 complexed with IP6 and BVM, independent of the cryo-EM analysis. To explore the energetics of BVM binding to CA_CTD_-SP1 ^47,48^, we calculated the free energy profiles using umbrella sampling^49^ simulations with Hamiltonian replica exchange^50^(US-HREX). This approach allowed us to sample BVM conformations along a reaction coordinate, focusing on the displacement of BVM within the central pore of the CA_CTD_-SP1 hexamer. We set up four separate systems with BVM in two different orientations—either with the dimethyl-succinate moiety up or down— and in the presence or absence of IP6. The resulting free energy profiles confirm kcal/mol) CA_CTD_ (up orientation) is energetically more favorable, regardless of the presence or absence of IP6 (**Fig. 5a**). The binding energy of BVM in the dimethyl-succinate up orientation (∼ -25 kcal/mol) approximately (∼ -12 kcal/mol). In addition, the BVM binding energy does not deviate significantly with or without IP6 in the six-helix bundle. Specifically, for the BVM-up orientation, we calculated a binding energy of -24.7 ± 0.1 kcal/mol in the presence of IP6 and -23.5 ± 0.3 kcal/mol in its absence. This is consistent with experimental predictions that BVM and IP6 binding are not competitive^19^.

**Fig. 5.**
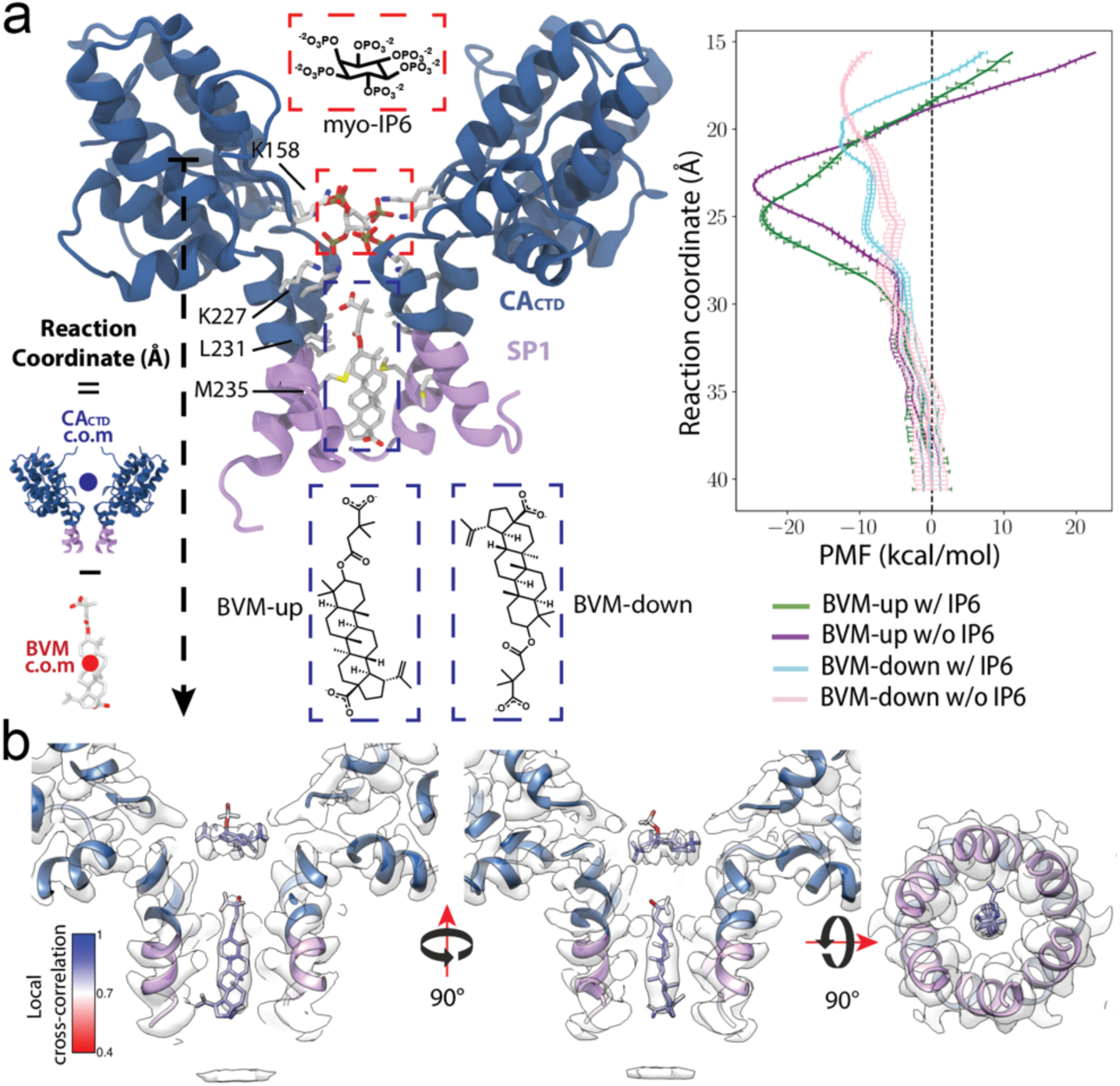
MD simulations of BVM binding. a. BVM binding characterized by potentials of mean force (PMF). Left: The binding site of BVM in the CA_CTD_-SP1 hexamer is probed by pulling BVM along a reaction coordinate defined as the z-axis projection of the distance between the center of masses (c.o.m.) of the CA_CTD_ and BVM. Right: The PMF for the interaction of BVM with CA_CTD_-SP1 is measured along the reaction coordinate for two possible orientations of BVM (dimethyl-succinate up or down) and in the presence or absence of myo-IP6. The PMF curves characterize the position and energy barrier for the binding of BVM in the CA_CTD_-SP1 6-helix bundle. CA_CTD_ is colored in dark blue and SP1 in lavender. b. BVM binding mode from MD simulations in relation to cryo-EM density. Front, side, and bottom views of the BVM binding pose at the energy minimum from the PMF after MDFF refinement. The BVM binding position from the PMF matches the observed cryo-EM density with high cross-correlation coefficients. CA_CTD_, BVM, and IP6 are colored by local cross-correlations according to the color bar. Cryo-EM density is shown in white.

In the energy minimum for the BVM-up orientation, the betulinic acid moiety is surrounded by hydrophobic residues L231 from CA_CTD_ and M4 from SP1, which can contribute to stabilizing bound BVM via methionine-π interactions. The simulations further indicate that BVM in the up-orientation is stabilized by the electrostatic interaction of dimethyl-succinate’s carboxylate group and the ɛ-amino group of K227 (**Fig. S4**). When BVM is far from the minimum in the energy profile, all six K227 residues in the six-helix bundle are oriented towards IP6. However, as BVM moves along the reaction coordinate towards the energetic minimum, some K227 residues are reoriented to form salt bridges with the dimethyl-succinate moiety. This is consistent with the cryo-EM density where the K227 has alternative conformations pointing towards either IP6 or BVM (Fig #).

Furthermore, the independently derived binding position of BVM in the CA_CTD_-SP1 six-helix bundle from the free energy profile calculations is consistent with the BVM density observed by cryo-EM. We extracted the position of BVM and IP6 from the energy minimum and refined their position in the Gag_CA-SP1_ hexamer density using molecular dynamics flexible fitting (MDFF)^50^. The position is closely aligned with the density, as measured by high local correlation values (**Fig. 5**). Notably, one of the IP6 phosphate groups shifts away from the central axial position upon BVM binding in the simulation (**Fig. 5b**), which is consistent with the cryo-EM density showing an altered binding pose of IP6 in the presence of BVM (**Fig. 4e**). Together, this analysis demonstrates that combining MD simulations with cryo-EM data from PFO-treated VLPs enables high-resolution *in situ* structural determination of inhibitors bound to their viral targets, providing new insights into BVM binding to the immature Gag_CA-SP1_ lattice that have previously been challenging to obtain.

## Discussion

Proper formation of the immature Gag lattice and subsequent capsid maturation are vital steps within the HIV-1 lifecycle. Inhibitors, such as Bevirimat (BVM) and Lenacapavir (LEN), have been known to alter the assembly and proper maturation of HIV-1 virions, but high-resolution structural information of their binding to the native immature lattice has yet to be defined. This led to a deficit in the mechanistic understanding of BVM and LEN binding, which is critical for the continued development and improvement of these inhibitors. We overcame this challenge by providing high-resolution descriptions of the binding modes of BVM and LEN within the native immature lattice of mammalian-produced virus-like particles (VLPs), utilizing in situ single-particle cryo-EM.

LEN is the first FDA-approved capsid inhibitor and has been shown to bind and alter the dynamics of the mature capsid in the early stages of HIV-1 life cycle. Remarkably, our *in situ* structure of the LEN-bound VLPs has unveiled a new mechanism of action of LEN during the late stages of HIV-1 budding and maturation. Our analysis revealed that LEN can interact with the immature Gag lattice by engaging three CA_NTD_ domains. The primary LEN-binding CA_NTD_ interacts with LEN via the canonical FG-binding pocket in a manner similar to the mature capsid, while two additional CA_NTD_ copies, uniquely arranged in the immature lattice, form new interaction interfaces. In the mature CA lattice, LEN is coordinated by the primary interacting CA_NTD_ and the CA_CTD_ of a second CA chain within the same hexamer. By contrast, in the immature Gag lattice, LEN engages three CA_NTD_ chains, two within a hexamer and one from a neighboring hexamer. This binding mode allows LEN to bind to the immature Gag lattice with a unique mechanism whereby only the CA_NTD_ interfaces are involved, independent of the CA_CTD_-SP1 layer of the immature Gag lattice. Consistently, our data showed that the binding of LEN alters the CA_NTD_-CA_NTD_ interactions and reorients the positioning of CA_NTD_ with respect to the fixed CA_CTD_-SP1 (Fig **3b**). This leads to a curious phenomenon: the CA_NTD_ layer undergoes independent rearrangement despite being connected to MA at its N-terminus and the fixed CA_CTD_ -SP1 at its C-terminus. This flexibility in the immature Gag lattice may be important for its mechanical stability, pliability, and maturation process. Interestingly, the critical R18 residue at the immature CA_NTD_-CA_NTD_ interface is also involved in LEN binding, linking R18 to both LEN interaction and lattice dynamics. Since R18 plays a pivotal role in maturation by coordinating the IP6 molecule within the mature CA hexamer, its sequestration by LEN may disrupt maturation. These findings suggest that LEN functions in the late stages of the HIV-1 lifecycle by disrupting conserved interactions within the immature lattice and trapping residues in conformations incompatible with maturation, thereby contributing to its inhibitory mechanism against HIV-1.

Our study also sheds light on the binding mechanism of the maturation inhibitor BVM. With the combination of high-resolution *in situ* cryo-EM studies and MD simulations, we obtained a more accurate binding pose of BVM. This binding pose has the succinyl moiety of BVM interacting with and reorganizing the K227 residues, which likely affects the IP6 binding dynamics; while the betulinic acid moiety is bound between residues L231 of CA and T8 of SP1. This binding mode also provided some explanation to the BVM resistant mutations and polymorphisms, including L231F/M (CA) and A1V, Q6H/A, V7A (SP1) that are within the range of residues at the BVM binding site. Compared to previous structural studies, we find that the position of BVM in our *in situ* cryo-EM studies with enveloped VLPs has shifted from earlier models. This may be attributed to differences in experimental methods and resolutions. The earlier MicroED and NMR studies utilized *in vitro*-assembled Gag truncations (CA_CTD_-SP1). It is possible that the differences in experimental conditions and constructs led to the observed shift in BVM binding. Our study provides an accurate binding pose in the native virus, which, in combination with previous research, enhances our understanding of BVM binding dynamics and offers insights for future structure-based drug design targeting maturation inhibitors.

Remarkably, the combination treatment of both BVM and LEN demonstrates that the two compounds are compatible and do not interfere with each other. Specifically, (1) the binding pose of either BVM or LEN remains identical in single- and double-treated VLPs; (2) the CA_NTD_ conformations, both in the apo form and that altered by LEN, show no further change by additional BVM binding; and (3) IP6 adopts the same BVM-altered binding pose in both the BVM-only and double-treated structures. These findings suggest that BVM and LEN bindings are independent events, with their mechanisms of inhibition not affecting each other, resulting in only additive effects. This points to the possibility that a combination therapy simultaneously targeting the FG pocket with a CA inhibitor and the CA-SP1 junction with a maturation inhibitor could be an attractive therapeutic option worthy of further exploration.

This work highlights the power of the *in situ* single-particle cryo-EM approach in achieving high-resolution reconstructions of the Gag_CA-SP1_ lattice, captured in complex with inhibitors and co-factors in their native viral environment. The capability of this technique opens new avenues for exploring the intricate interactions between viral proteins and their binding partners, offering deep insights that can drive structure-based drug design and optimization of viral inhibitors. The knowledge gained from these studies will not only facilitate the design of more effective Gag-targeting inhibitors but also expand the range of high-resolution techniques available for studying large viral protein assemblies and their interactions with co-factors and therapeutic agents.

## Acknowledgements

This work was funded in part by the National Institutes of Health through grants T32GM008283, U54AI170791, and R01AI178846. Cryo-EM data was collected with the support of staff at the Brookhaven National Laboratory – Laboratory for BioMolecular Structure (LBMS) and the Yale CryoEM Resource. LBMS is supported by the DOE Office of Biological and Environmental Research (KP1607011). We thank the Xiong lab members for discussions.

## Methods

### VLP Production

Expi293F cells (Thermo Fisher Scientific) were cultured in 8% CO_2_ at 37°C while shaking at 250 rpm. VLPs were produced by transfection of Expi293F cells with 1 µg/mL NL-MA/NC DNA^35^ using the Expi293 Transfection Kit (Thermo Fisher Scientific). After 48 hours of expression, suspensions of VLPs and cells were subjected to centrifugation at 500 x g for 5 minutes to pellet the cells, and the supernatant was filtered through a 0.45 µm syringe filter. The filtered supernatant was then pelleted through a 20% sucrose cushion by ultracentrifugation in a Beckman rotor SW32 at 120,000 x g for 3 hours at 4°C. The resulting pellet was thoroughly resuspended in 600 µL 10 mM Tris-HCl, 100 mM NaCl, 1 mM EDTA, pH 7.4 (1X STE) before overlaying onto a 400 µL 15% sucrose layer over an 11 mL gradient of 30 to 70% sucrose that was poured over a 3 mL 85% sucrose cushion. This gradient was subjected to ultracentrifugation in a Beckman rotor SW27 at 100,000 x g for 16 hours at 4°C. The gradient was collected from the top down in 1 mL fractions. VLPs were typically recovered in fractions 3 to 5. The fractions containing VLPs were then diluted 2-fold in 1X STE and then subjected to centrifugation for 30 minutes at 16,000 x g at 4°C to concentrate the VLPs. The resulting pellets were resuspended in 1X STE and 10% sucrose to bring the VLPs to their desired concentration. VLPs were then aliquoted and frozen to -80°C for future use.

T8I VLPs were produced as previously described^7^. Briefly, HEK293T cells (ATCC CRL-3216) were transfected with a codon-optimized Gag expression plasmid (pCMV-Gag-opt)^51^. After 40 hours of expression, the culture supernatants were filtered through a 0.45 µm syringe filter and VLPs were pelleted through a 20% sucrose cushion by ultracentrifugation. The resulting pellet was resuspended in 1X STE buffer, aliquoted, and frozen to -80°C for future use.

### Perfringolysin O (PFO) Purification

The cytolytic toxin PFO gene from *Clostridium perfringens* was cloned as 6x His-tagged PFO constructs into a pET-21-derived vector (Novagen) and expressed in BL21 (DE3) E. Coli. Cells were grown in Terrific Broth (TB) medium at 37°C in a shaking incubator until they reached an OD600 of 0.8. Protein expression was then induced with 0.5 mM IPTG for 16 hours at 18°C before collection by centrifugation. Cell pellets were stored at -80°C until used. Then, cell pellets were resuspended in lysis buffer (50 mM Tris, pH7.5, 0.5 mM TCEP, 300 mM NaCl) with 1X HALT protease inhibitor (Thermo Fisher Scientific) and lysed using a microfluidizer. Whole-cell lysates were spun down to remove cell debris, and the supernatant of lysate was collected and filtered through 0.45 um filters and subsequently loaded into a 5mL HisTrap column (GE) on an AKTA system (GE) at a flow rate of 1 mL/min. The column was washed with lysis buffer and then eluted on a gradient of elution buffer containing an additional 500 mM imidazole. Purified PFO concentration was determined by Nanodrop (Thermo Fisher Scientific). The final PFO sample at a concentration of 10 mg/ml had 15% glycerol added, was flash-frozen in liquid nitrogen, and stored at -80°C until used.

### Perforated VLP sample preparation

Purified VLPs were pelleted by centrifugation for 30 minutes at 16,000 x g at 4°C to remove any residual sucrose and resuspended in 1X STE. The VLPs are enriched by spinning down and resuspending in the same buffer. The final estimate of Gag concentration is about 50-150 uM, determined by p24 quantification. The resulting VLPs were then mixed with various desired components: PFO (final concentration 2 µM), IP6 (final concentration 4 mM), BVM (final concentration 100 µM), and LEN (final concentration 300 µM) and incubated at 37°C for 5-15 minutes. Perforated VLPs are immediately used for cryo-EM sample preparation.

### Negative stain EM

Grids for negative stain EM analysis were prepared by applying 4 µL of sample on a glow-discharged Carbon Film 400 mesh Cu grid (Electron Microscopy Sciences), manually blotting with a filter paper, and staining with 2% uranyl acetate for 1 minute. The grids were then imaged using a Talos L120C microscope (Thermo Fisher Scientific).

### Cryo-EM Imaging

Grids for cryo-EM were prepared by applying 4 µL of sample on a glow-discharged Quantifoil 300 mesh Cu 2/1 grid (Quantifoil Micro Tools), blotting for 7.5 seconds, then plunge-freezing in liquid ethane using a Mark IV FEI Vitrobot (Thermo Fisher Scientific). The grids were then screened, and data were collected on a 200 keV Glacios electron microscope (Thermo Fisher Scientific) with a K3 direct detection camera (Gatan) with a magnification of 45000x, physical pixel size of 0.86 Å and total dose of 50 e^-^/Å^2^. The final data were collected on a 300 keV FEI Titan Krios (Thermo Fisher Scientific) with a K3 direct detection camera (Gatan) and energy filter (Thermo Fisher Scientific) at the Yale cryo-EM Resources facility or the Brookhaven National Laboratory LBMS facility with a magnification of 81000x, physical pixel size of 1.06 Å, energy filter slit at 20 eV and total dose of 50 e^-^/Å^2^.

### Cryo-EM Data Processing

Data processing of cryo-EM images was done using CryoSPARC v4.4.1 (**Table 1**)^52^. Raw movies were motion and CTF corrected using the implementation in CryoSPARC. Initial particles were manually picked and followed by 2D classification to generate templates for template particle picking^52^. Side view and top view classes were used in separate templated particle picking jobs for efficient picking of both views. For the data sets with a pixel size of 1.06 Å collected on the Krios microscope, the initial particles were extracted with a box size of 282 pixels and Fourier cropped to 64 pixels. To remove junk particles and perform initial 3D alignment to the particles, iterative cycles of heterogeneous refinement were performed with reference volumes including a previously resolved map of immature CA lattice (EMD-13354)^53^ and 3 copies of a featureless flat-ellipsoid-shaped decoy map, generated from the reference volume low-path filtered to 200 Å resolution. The featureless map is for absorbing junk particles into the junk classes. C6 symmetry is implemented in the heterogeneous refinements. Upon convergence after a few rounds of heterogeneous refinement, the well-defined class of particles was re-extracted with a box size of 282 pixels without cropping. The re-extracted particles were then refined with local refinement. 3D classification and subsequent local refinements were performed to select the final well-defined particle set. The final particle set then underwent global and local CTF refinements^54^ and reference-based motion correction, followed by a final round of local refinement. The local resolution range was analyzed by a local resolution job (**Table 1**). The detailed processing steps for each data set are illustrated in supplemental figures (**Fig. S8, S9, S10**).

### Model building and refinement

Atomic models were built for the T8I intact data by docking PDB 7ASL^7^, into the map using ChimeraX^55^, followed by iterative manual rebuilding in Coot^56^ and real-space refinement using Phenix (phenix.real_space_refine)^57^. For accurate model building, the maps were sharpened by phenix anisotropic sharpening and converted to mtz format by CCPEM^58,59^. The map symmetry parameters of C6 were determined by phenix.map_symmetry and were used during the real space refinement and applied to the final model by phenix.apply_ncs implemented in Phenix^60^. The ligands including IP6, LEN, and BVM and their parameters were generated using phenix.elbow^61^ and imported into Coot as a CIF dictionary file for geometry restraints. The figures were made in ChimeraX^55^ and PyMol (Schrödinger, LLC).

### Molecular dynamics system preparation

In preparation for performing molecular dynamics simulations of the CA_CTD_-SP1-BVM complex, we built our system from the NMR atomic coordinates of CA_CTD_-SP1 in complex with IP6 and BVM (PDBID 7R7Q and 7R7P, respectively)^62^. Hydrogens were added to the protein according to the predicted protonation states of each amino acid at pH 7.0 using the propKa3 software^63^. The CA_CTD_-SP1 model in complex with BVM was then solvated with TIP3P water molecules^64^ into a P6 hexagonal unit cell with director vectors of length 70.94 Å, 70.94 Å, and 110.25 Å using the VMD *solvate* plugin^65^ and in-house TCL scripts. The solvated systems were ionized to 150 mM NaCl and charge neutrality using the VMD *autoionize* plugin^65^. Following this procedure, we prepared two systems of CA_CTD_-SP1 in complex with BVM in the presence or absence of IP6. Each prepared system encompassed over 49,000 atoms.

The prepared systems were then minimized and equilibrated by the following procedure. First, the solvent in each system was minimized for 5000 steps using the gradient descent algorithm while keeping the protein, IP6, and BVM molecules fixed. A second minimization step was then performed allowing the IP6 and BVM molecules to move and restraining the protein backbone atoms with harmonic constraints with a force constant of 10 kcal/mol Å. Each minimization step was followed by a gradual heating step where the temperature of the system was raised from 50K to 310K at a rate of 20 K/ns with the same restraints as the minimization. After tempering, the protein harmonic restraints were gradually removed at a rate of -1 kcal/mol Å every 0.2 ns in an NPT simulation at T = 310 K and P = 1 atm. Finally, each system was equilibrated in an NPT ensemble at T = 310 K and P = 1 atm for 100ns. Once equilibrated, the last snapshot of each equilibration simulation was used as the initial structure of any further production simulations.

All MD simulations were performed in the NAMD simulation engine, minimizations were performed in NAMD 2.14^66^, and all other steps were performed in NAMD3 alpha 7 ^67^. The force field parameters for proteins, water molecules, and ions were extracted from the CHARMM force field version c36^68–71^ while parameters for IP6 were generated from analogy to the CHARMM force field for small molecules using CGenFF^72^. CGenFF-generated parameters for BVM had high penalties. Therefore, BVM was parameterized from QM-level calculations as detailed in the supplementary information. During MD simulations, the temperature was controlled using the Langevin thermostat with a damping coefficient of 0.1 ps^-1^. In NPT simulations, the pressure control was applied via the Nose-Hoover Langevin piston method with a period of 200 ps and a decay time of 100 ps. Periodic boundary conditions were enforced, and pressure control maintained the area of the XY plane of the unit cell, allowing for fluctuations in the Z axis. All bonds to hydrogen atoms were constrained with the SHAKE^73^ and SETTLE^74^ algorithms. Nonbonded interactions were calculated with a cutoff distance of 12 Å with the long-range electrostatic interactions were calculated every 4.0 fs using the Particle-Mesh-Ewald^75^ summation with a grid size of 1.0 Å.

### Molecular mechanics parameterization of BVM

The reliability of molecular dynamics simulations to investigate protein-ligand dynamics and energetics depends on the molecular mechanics force field used to describe the intra and inter molecular potentials. Thus, a force field that accurately represents the molecular mechanics parameters of the ligands is crucial for our simulations. Our previous work described in detail the parameterization process for obtaining CHARMM molecular mechanics forcefield parameters for IP6 and BVM^19^. In brief, initial force field parameters for IP6 and BVM were derived by analogy to the CHARMM general forcefield for small molecules using CGenFF version 2.5.1 ^72^. All CGenFF parameters for IP6 had penalties lower than ten, indicating good analogy, and were thus used directly in the simulations. BVM CGenFF parameters with penalties greater than 10 however were refined at the QM level using the Force Field ToolKit v2.1^76^ (FFTK) plugin in VMD^65^ 1.9.4 and Gaussian16^77^. In all simulations presented, we utilized the QM-optimized BVM parameters.

To probe the BVM QM-optimized force-field parameters and additionally to analyze the dynamics of BVM, we performed 100ns molecular dynamics simulations of BVM bound to Gag_CTD-SP1_ and isolated BVM in explicit water utilizing the CGenFF force field parameters and the QM-optimized parameters for comparison (**Fig. S5**).

Atomic charges predicted by analogy via CGenFF had penalty values in the 10-15 range and were optimized at QM level to reproduce hydrogen bond interactions with a water molecule. As shown in Figure S5, after optimization, most atomic charges have little variation from the CGenFF values (Δ*q*∼10^−2^), with the greatest variations being present in the charge of terminal carbons in the dimethyl-succinate moiety (Δ*q* ∈ (0.06, 0.18)).

Analyzing the trajectories of isolated BVM and in complex with Gag_CTD-SP1_ we observe no discrepancy in the root mean square fluctuations (RMSF) for BVM atoms measured over 100ns using CGenFF and QM-optimized parameters. In both cases, the betulinic acid backbone is relatively rigid, while the dimethyl-succinate moiety is flexible (**Fig. S5b,c**) and plays a crucial role in interacting with the Lys227 ring inside the six-helix bundle of Gag_CTD-SP1_.

Although the observed fluctuation of atomic positions is consistent for BVM using CGenFF or QM-optimized parameters, the probability of the conformational states the dimethyl-succinate moiety adopts during MD sampling is considerably different for CGenFF and QM-optimized parameters. We characterized the rotation of the dimethyl-succinate moiety by measuring the dihedral torsion angle of the carbon atoms in this moiety throughout the MD simulation and identified three meta-stable states (A, B and C) with different populations. As observed in **Figure S5d**, in isolated BVM state B has a higher probability when using the CGenFF parameters, while state C has the higher probability for the QM-optimized parameters. Furthermore, during the BVM bound to Gag_CTD-SP1_ simulations, we observed the dimethyl-succinate moiety visiting the three metastable states with the QM-optimized parameters, while sampling with the unoptimized CGenFF parameters results in state A not being occupied in the 100ns trajectory.

Overall, our results show that although there is no significative difference in the charge and dynamics between the CGenFF and QM-optimized for the betulinic acid moiety due to the rigidity of pentacyclic triterpenoids, the QM-optimized parameters have a major impact on the conformational space available for the dimethyl-succinate moiety which is critical for the interaction of BVM in the Gag_CTD-SP1_ six helix bundle.

### Potentials of mean force (PMF) via Umbrella sampling Hamiltonian replica exchange (US-HREX)

Free energy profiles of BVM binding to Gag_CTD-SP1_ hexamer were obtained by performing Umbrella Sampling^48^ (US) simulations with Hamiltonian Replica Exchange^49^ (HREX). For these simulations, we defined a reaction coordinate as the z-axis projection of the displacement vector between the center of mass of the Gag_CTD-SP1_ hexamer (*r⃗_CTD-SP_*_1_) and the center of mass of the BVM (*r⃗_BVM_*) molecule

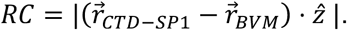

This reaction coordinate allows us to track the displacement of BVM along the axis of the central pore of the Gag_CTD-SP1_ hexamer. The initial configurations for each US simulation were generated by pulling BVM along the reaction coordinate at a rate of 0.75 Å every 2 ns in steered molecular dynamics (SMD) ^78,79^ simulations. A total of 64 snapshots from the SMD trajectory were saved every 2 ns and utilized as the initial coordinates for each US window. The biasing potential *U_i_*(*X*) for the US window *i* = 1, … ,64 is set to a harmonic potential calculated on the reaction coordinate for conformation X:

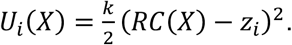

With spring constant 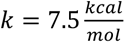 Å and harmonic potential center *z_i_* covering the range of *z* = 10 Å to *z* = 45 Å with a uniform spacing of 0.75 Å. To enhance the sampling efficiency and facilitate the convergence of the US simulations, we utilized Hamiltonian Replica Exchange (HREX) molecular dynamics. In HREX, each window in the US simulation is a replica that can exchange conformations with a neighboring replica with a given transition probability. In our simulations, a replica exchange is attempted every 2 ps and it is accepted with the swap probability between neighboring replicas *i* and *j* is given by the Metropolis criterion^49^ 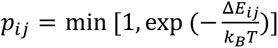. Where *T* = 310*K* is the simulation temperature, *k_B_* is the Boltzmann constant and Δ*E_i,j_* = *K*(*RC_i_* − *RC_j_*)(*z_i_* − *z_j_*) is the energy difference between neighboring replicas *i* and *j*.

After performing the US-HREX simulations, the biased probability from US at each window was analyzed via the weighted histogram analysis method (WHAM) ^80,81^ to compute the potentials of mean force (PMFs). US-HREX simulations were performed for 50 ns for each system, and data from the first 25 ns was removed before PMF calculations. The convergence of the PMF was evaluated by confirming no significant changes in the energy profile as the sampling time increased.

### Molecular Dynamics Flexible Fitting

We refined the BVM position in the Gag_CTD-SP1_ hexamer pore by performing molecular dynamics flexible fitting (MDFF)^50^ guided by the cryo-EM density. From the PMFs, we extracted the coordinates of the CA_CTD_-SP1 hexamer, IP6 and BVM molecules and aligned them to the density by performing rigid-body fitting of the CA_CTD_-SP1 domains. The BVM coordinates were then located in the T8I CA_CTD_-SP1 structure modeled from the density and further refined via MDFF by the following procedure: First the system was minimized via for 35000 steps using the conjugated gradient descent scheme ensuring convergence to values below 10 *kcal* ⋅ *mol*^−1^ ⋅ Å^−2^. Afterwards we performed a short molecular dynamics equilibration for 1ns in the NPT ensemble, maintaining the temperature at T=310 K using a Langevin thermostat with a damping coefficient of 0.1 ps^-1^ and pressure at P=1 atm via a Nose-Hoover Langevin piston with a period of 200 ps and a decay time of 100 ps, keeping the unit call area constant in the XY plane and allowing fluctuations in the Z axis. During both minimization and equilibration, we applied the cryo-EM density as a biasing potential coupled to the heavy atoms using the gridForces module in NAMD 2.14^66^ with a coupling grid scaling factor of 0.3. The fitting of the BVM and IP6 coordinates after MDFF were then evaluated by calculating the local cross-correlation coefficients using ChimeraX’s vop command^46^.

