## Supplementary material for "Structural insights into inhibitor mechanisms on immature HIV-1 Gag lattice revealed by high-resolution *in situ* single-particle cryo-EM": Suppliemental Figures

**Supplementary information:**

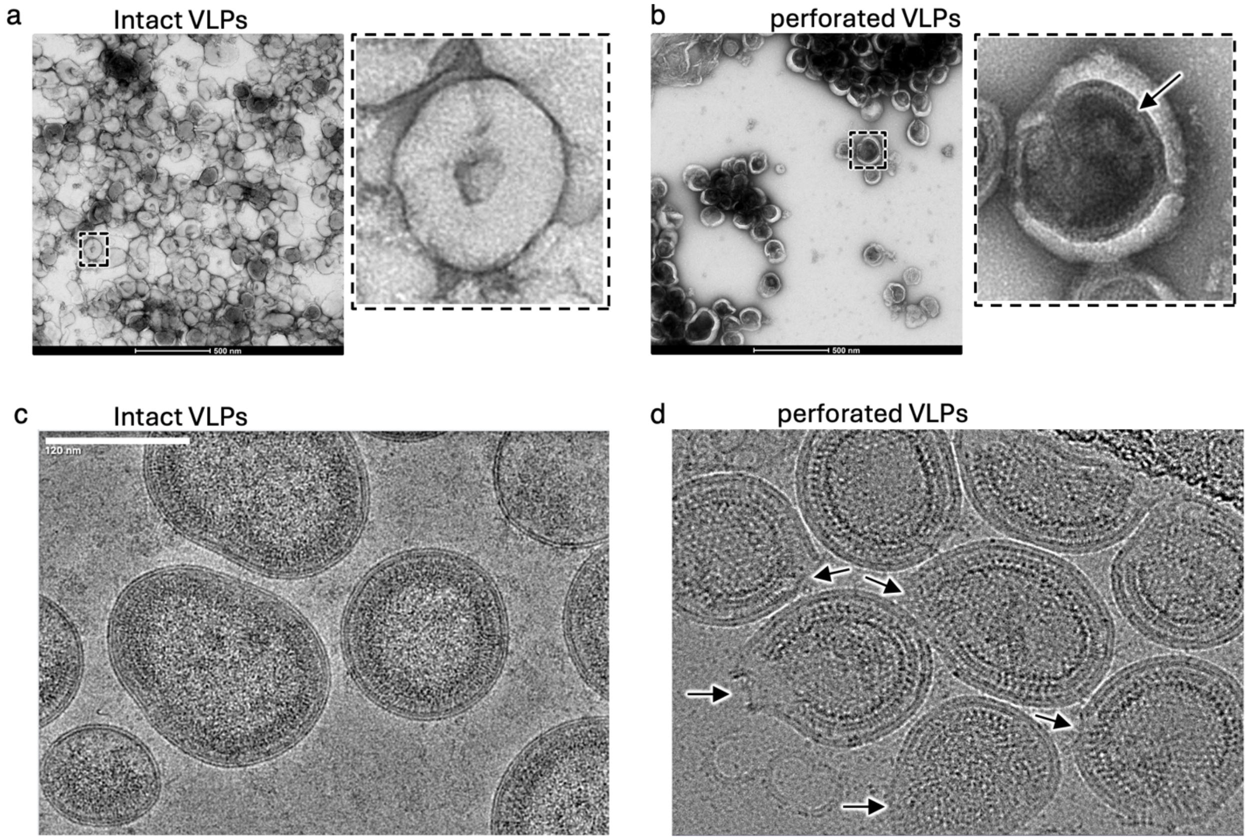


**Fig. S1 | PFO treatment to generate perforated VLPs**

a. Negative stain EM micrograph of intact immature VLPs.
b. Negative stain EM micrograph of PFO-perforated immature VLPs. The black arrow marks the immature Gag lattice, made visible as the negative stain enters the VLPs through the PFO-mediated pores.
c. Cryo-EM micrograph of intact immature VLPs.
d. Cryo-EM micrograph of PFO-perforated immature VLPs. The black arrows mark PFO-mediated pores.

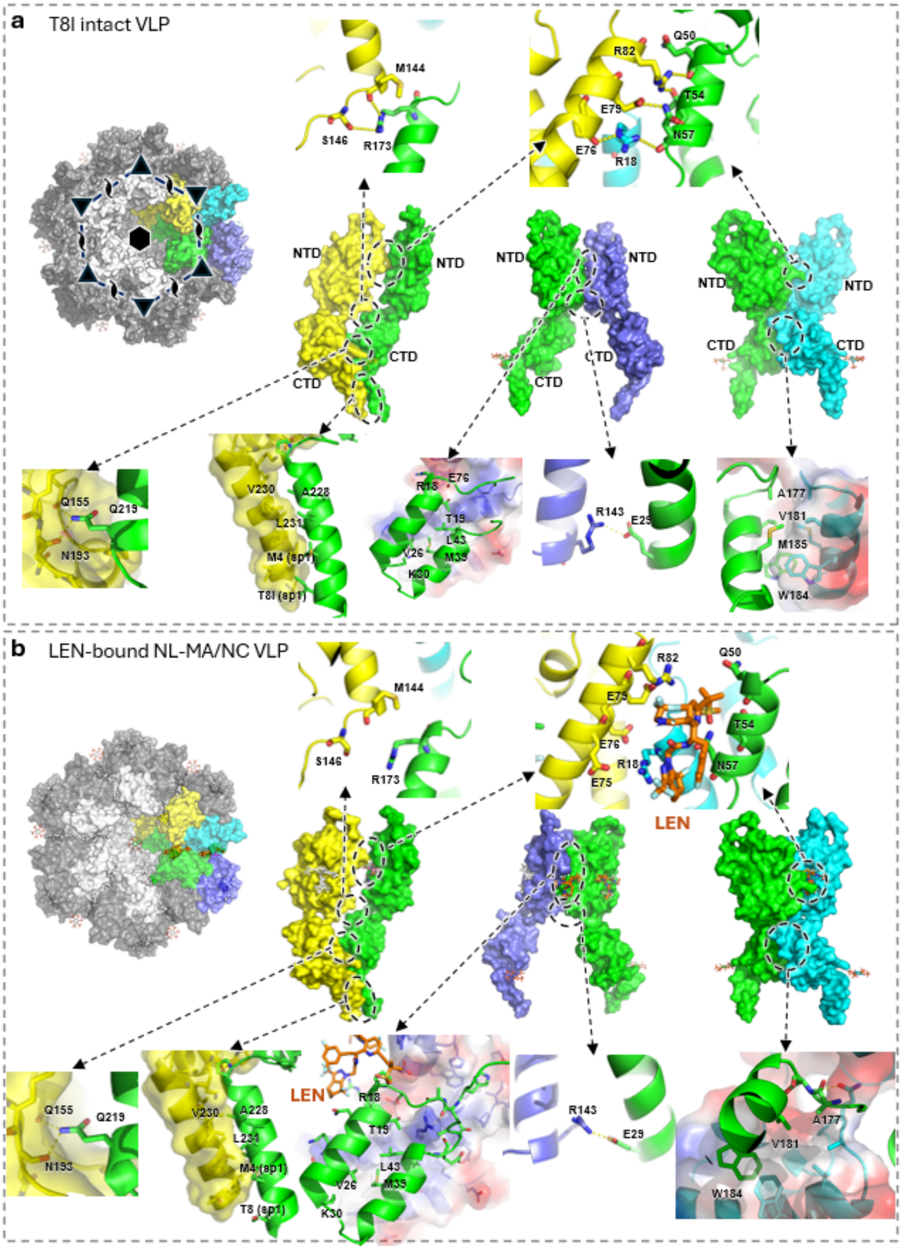


**Fig. S2 | Major interfaces in the immature Gag$_{CA\text{-}SP1}$ lattice from intact VLPs and LEN-treated VLPs.**

a. Major interfaces in the Gag$_{CA\text{-}SP1}$ lattice from intact T8I VLPs. Two CA-SP1 protomers in the same immature Gag$_{CA\text{-}SP1}$ hexamer are colored in green and yellow, while two CA-SP1 protomers from an adjacent hexamer are in cyan and purple.

b. Major interfaces in the Gag$_{CA\text{-}SP1}$ lattice from perforated LEN-bound NL-MA/NC VLPs. Two CA-SP1 protomers in the same immature Gag$_{CA\text{-}SP1}$ hexamer are colored in green and yellow, while two CA-SP1 protomers from an adjacent hexamer are in cyan and purple. The LEN molecules bound are shown in orange.

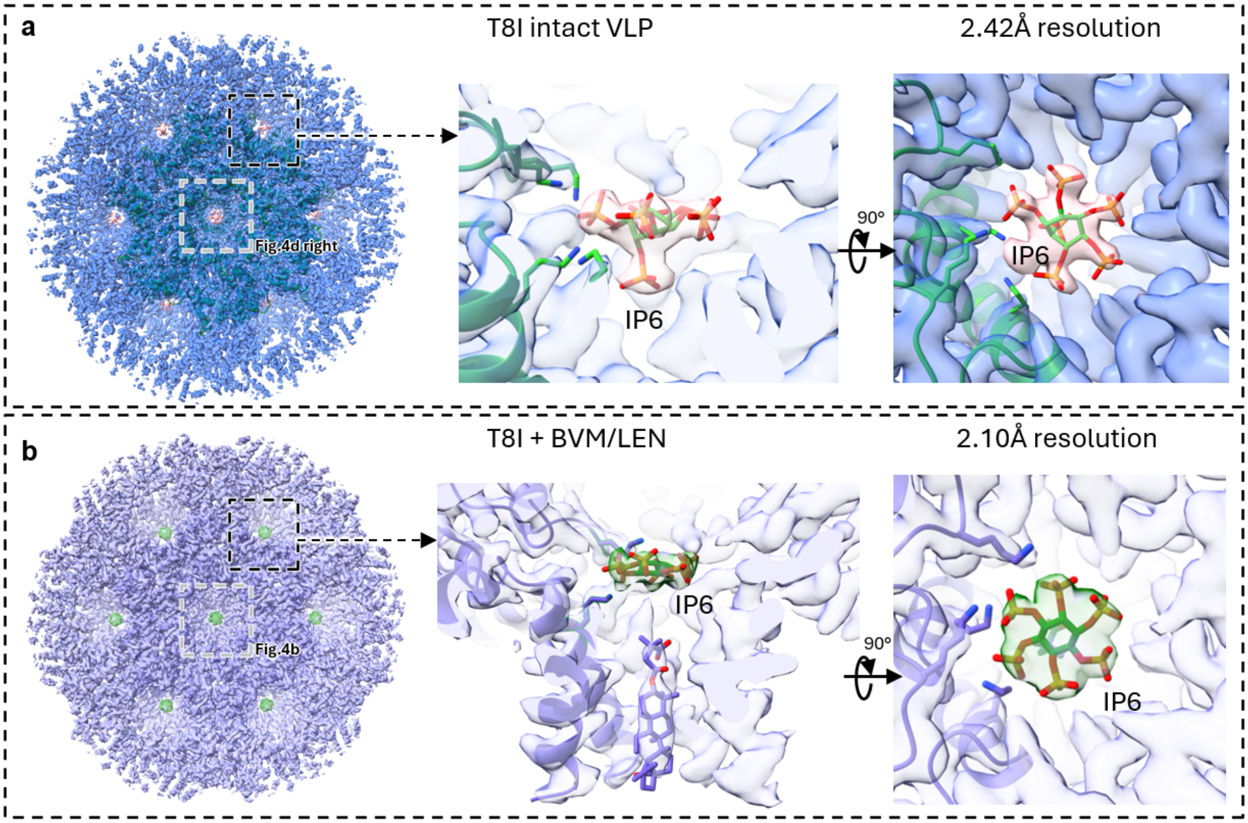


**Fig. S3 | IP6 binding sites of the peripheral hexamers in cryo-EM reconstructions of the immature $Gag_{CA\text{-}SP1}$ lattice.**

a. Cryo-EM map of the immature $Gag_{CA\text{-}SP1}$ lattice assembly from T8I intact VLPs is shown in blue. The inset shows side and top views of IP6 (orange) within the CA-SP1 hexamer pore from a non-central hexamer, boxed in black. The central hexamer shown in main Fig. 4d is boxed in grey.

b. Cryo-EM map of the immature $Gag_{CA\text{-}SP1}$ lattice assembly from T8I VLPs treated with BVM and LEN is shown in slate. The inset shows side and top views of IP6 (green) and BVM (slate) within the CA-SP1 hexamer pore from a non-central hexamer, boxed in black. The central hexamer shown in main Fig. 4d is boxed in grey.

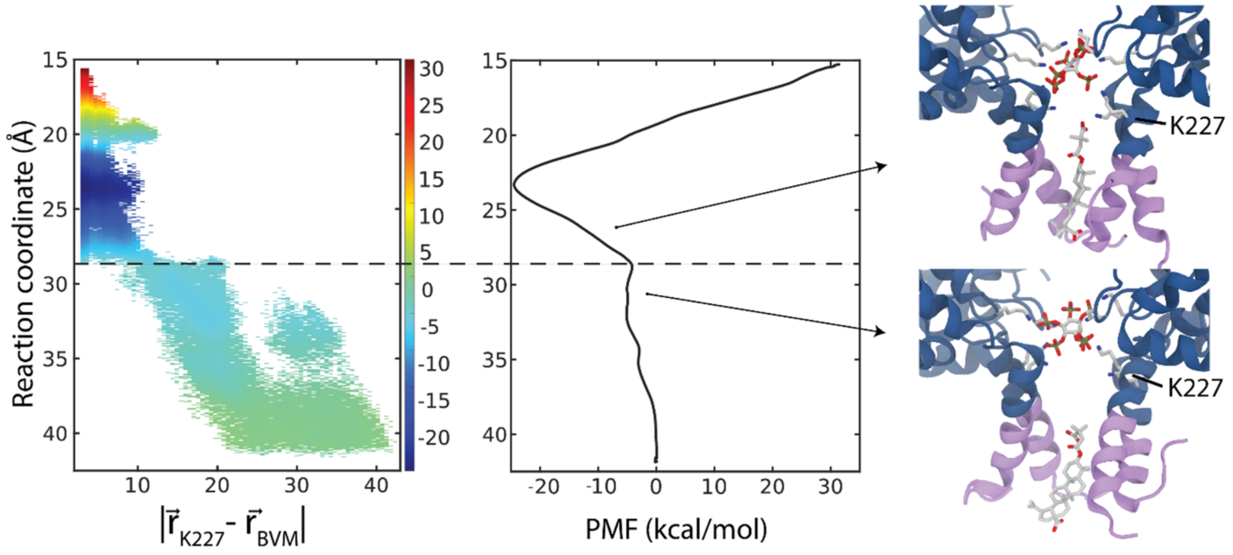


**Fig. S4 | Energetics of BVM interaction with K227 ring in the Gag$_{CTD-SP1}$ six helix bundle.** (Left) Correlation between BVM free energy of binding and the distance between BVM's dimethyl-succinate moiety and the amino group of K227 as measured via joint equilibrium probability[52]. The dashed line indicates the energy barrier for the free energy profile (Center). (Right) Two snapshots of BVM dynamics in the Gag$_{CTD-SP1}$ along the reaction coordinate (RC). Near the energy minimum, a salt bridge interaction forms between BVM and K227. Upon passing the energy barrier (RC >29 Å) the salt bridge is broken and BVM dissociates.

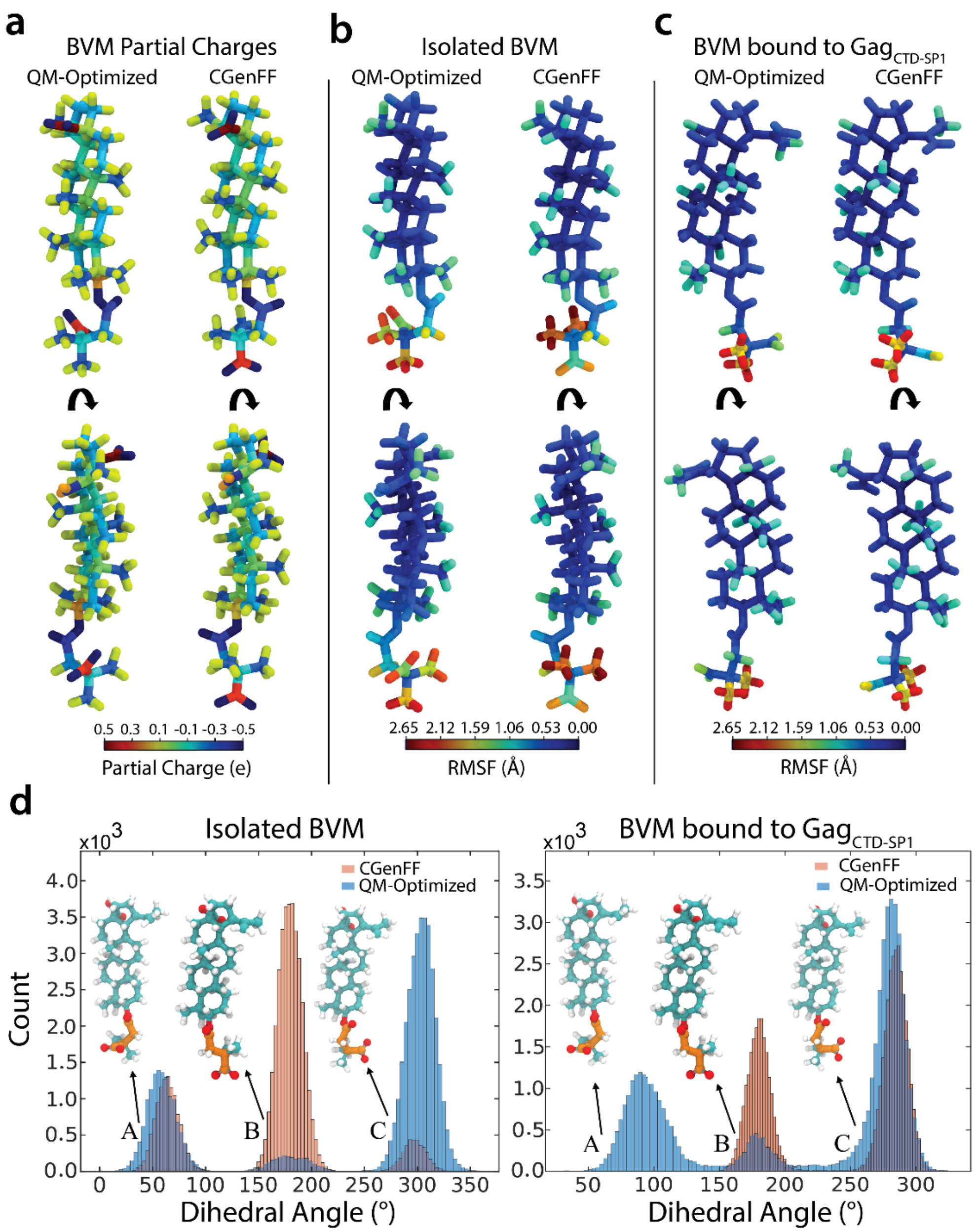


**Fig. S5 | Comparison between CGenFF and QM-Optimized BVM force field parameters.**

a. BVM atomic charge CGenFF assignment and QM-Optimized charges.
b. Per atom Root Mean Square Fluctuation (RMSF) from MD simulations of BVM in explicit water solvent.
c. Per atom RMSF from MD simulations of BVM in complex with Gag$_{CTD\text{-}SP1}$ hexamer in the presence of IP6.
d. Dihedral torsion angle conformational states of the dimethyl-succinate moiety as measured from the MD simulations of isolated BVM or in complex with Gag$_{CTD\text{-}SP1}$ hexamer in the presence of IP6. The probed dihedral is colored orange. BVM is colored by atom names in CPK representation.

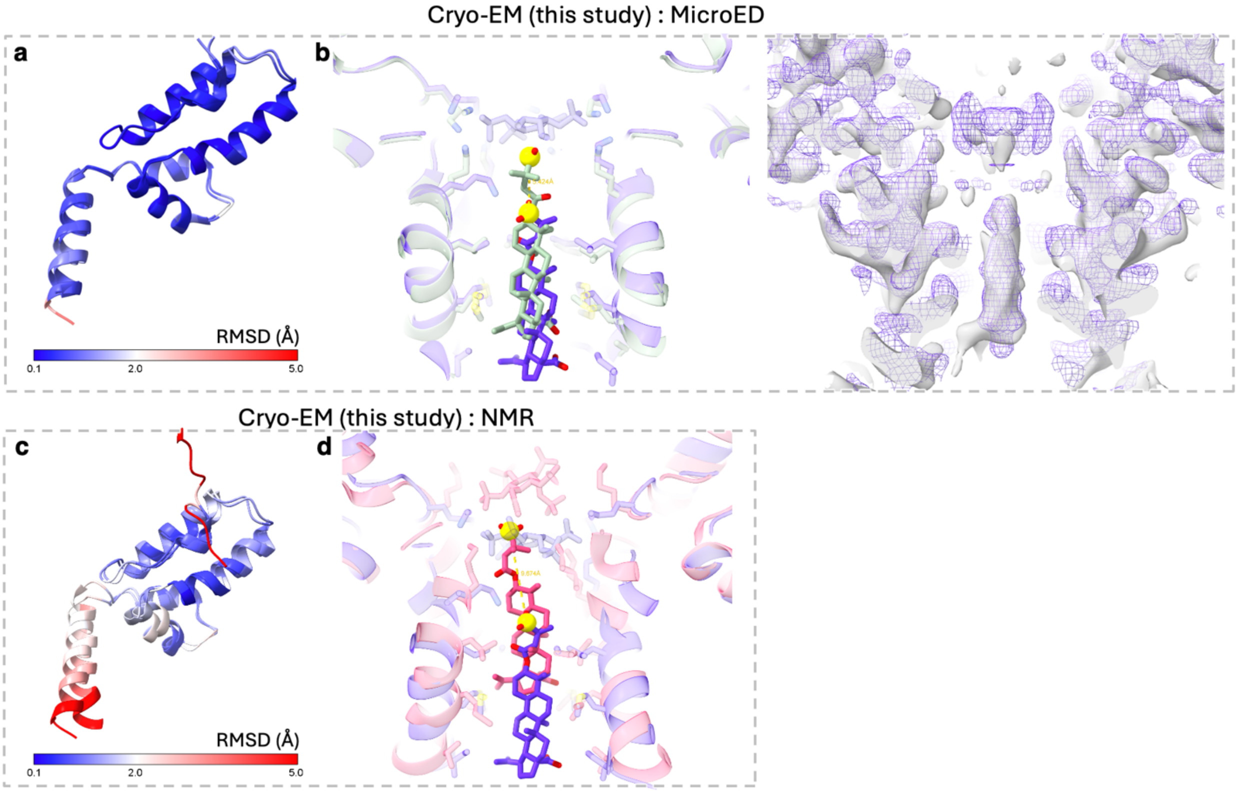


**Fig. S6 | Structural comparison of BVM-bound models from Gag lattice in VLPs (this *in situ* study) and previous *in vitro* assembled $CA_{CTD}$-SP1 lattice**

**a**. The $Gag_{CA\text{-}SP1}$ model from LEN/BVM-bound T8I VLPs (this study) is aligned with the previously reported MicroED structure of $CA_{CTD}$-SP1 (PDB 6N3U). The RMSD between 91 aligned C-alpha atom pairs is 0.5 Å.

**b.** Left: The $Gag_{CA\text{-}SP1}$ model from LEN/BVM-bound T8I VLPs (this study, slate) is superimposed with the microED model (6N3U, light lime). Because the 6N3U PDB entry does not include a BVM model, a BVM structure was reproduced based on the original microED density (EMD-0337) and the corresponding report[18]. The distance between the two BVM molecules from each model is around 5 Å, which is measured by the location of the succinyl group (marked by a yellow sphere). Right: A comparison of the cryo-EM map from LEN/BVM-bound T8I VLPs (this study, slate mesh) with the previously reported microED density map (EMD-0337, white surface). Except for the region corresponding to the IP6-binding site, most of the density is consistent between the two maps.

**c.** The $Gag_{CA\text{-}SP1}$ model from LEN/BVM-bound T8I VLPs (this study) is aligned with the previously reported NMR structure of $CA_{CTD}$-SP1 (PDB 7R7P). The RMSD between all 97 C-alpha atom pairs is ~3.5 Å. The helix at the CA-SP1 junction has the largest difference between the two models.

**d.** The $Gag_{CA\text{-}SP1}$ model from LEN/BVM-bound T8I VLPs (this study, slate) is superimposed with the NMR model (PDB 7R7P, pink). The distance between the two BVM molecules from each model is around 10 Å, which is measured by the location of the succinyl group (marked by a yellow sphere).

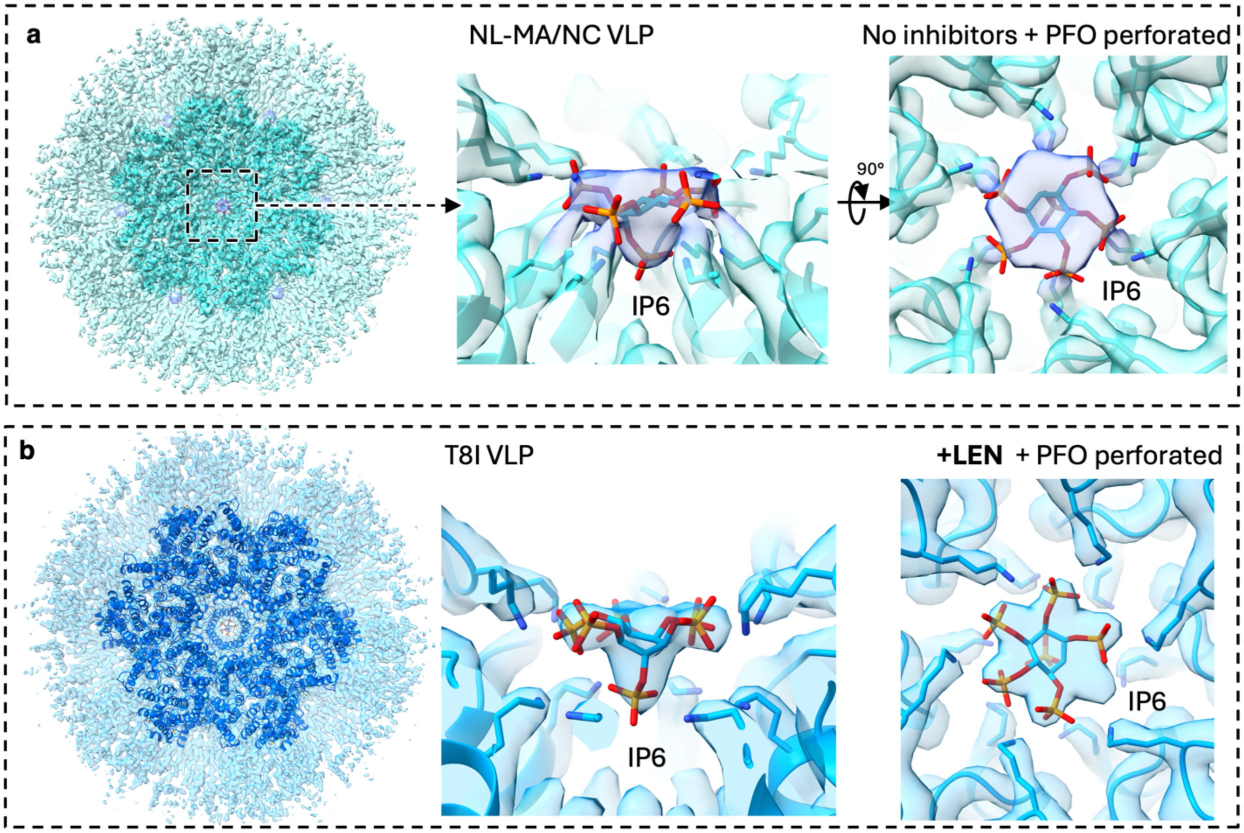


**Fig. S7 | LEN treatment does not alter the binding pose of IP6.**

- **a.** Cryo-EM map of the immature $Gag_{CA\text{-}SP1}$ lattice from perforated NL-MA/NC VLPs without treatment of inhibitors. The middle and right panels show side and top views of IP6 within the pore of CA-SP1 6-helix bundle in the central hexamer.
- **b.** Cryo-EM map of the immature $Gag_{CA\text{-}SP1}$ lattice from perforated and LEN treated T8I VLPs. The middle and right panels show side and top views of IP6 (blue) within the pore of CA-SP1 6-helix bundle in the central hexamer. The IP6 pose is not changed in the LEN treated VLP samples.

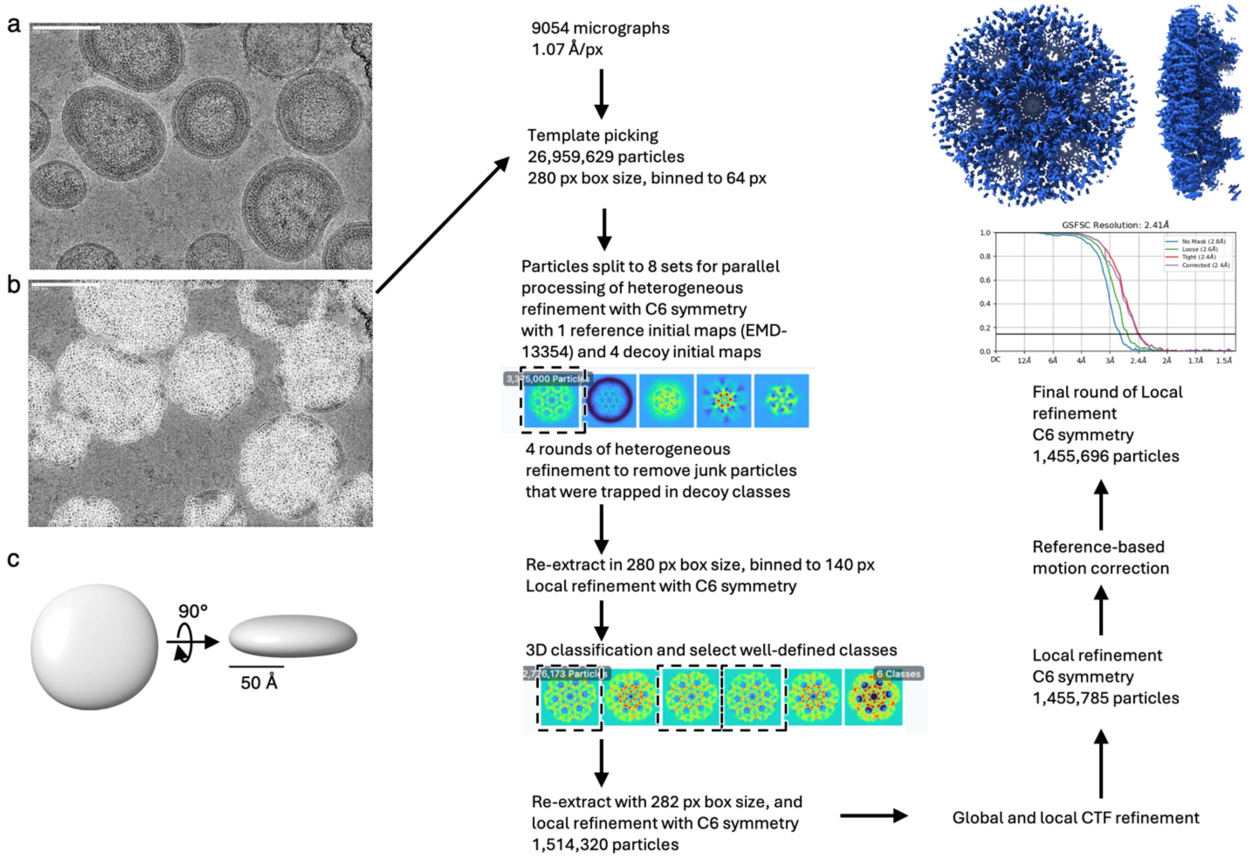


**Fig. S8 | Cryo-EM workflow for intact T8I VLPs.**

a. An example cryo-EM micrograph after motion correction.

b. Templated particle picking on the example micrograph and data processing workflow for the reconstruction of the immature $Gag_{CA\text{-}SP1}$ from intact T8I VLPs.

c. A featureless decoy map generated from the reference volume (EMD-13354), low-path filtered to 200 Å resolution, for trapping junk particles in heterogeneous refinement.

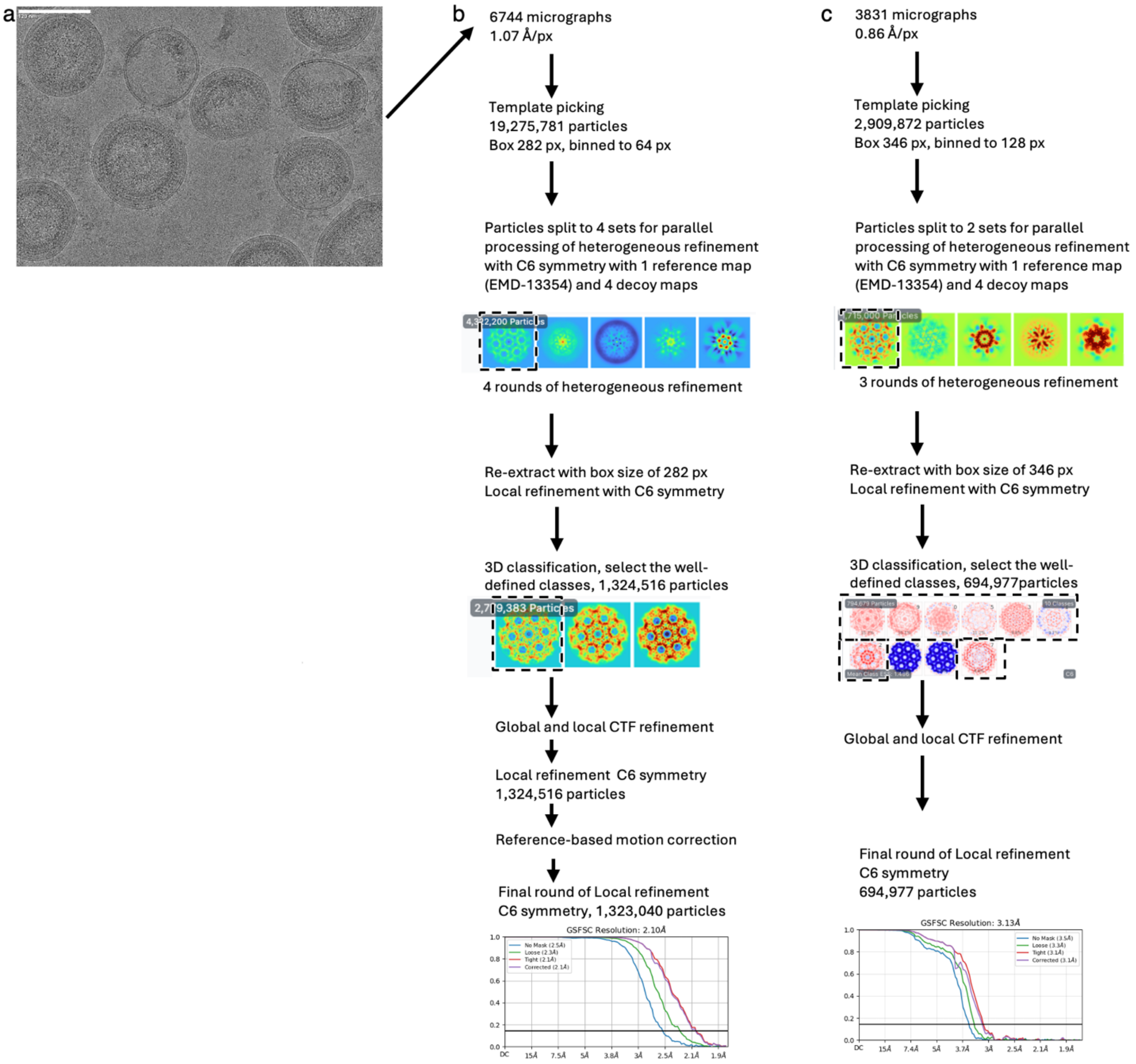


**Fig. S9 | Cryo-EM workflow for PFO-perforated BVM and LEN-bound T8I VLPs.**

a. An example cryo-EM micrograph after motion correction.

b. Data processing workflow for the reconstruction of the immature $Gag_{CA\text{-}SP1}$ with BVM- and LEN-bound PFO-perforated T8I VLPs.

c. Data processing workflow for the reconstruction of the immature $Gag_{CA\text{-}SP1}$ with LEN bound PFO-perforated T8I VLPs.

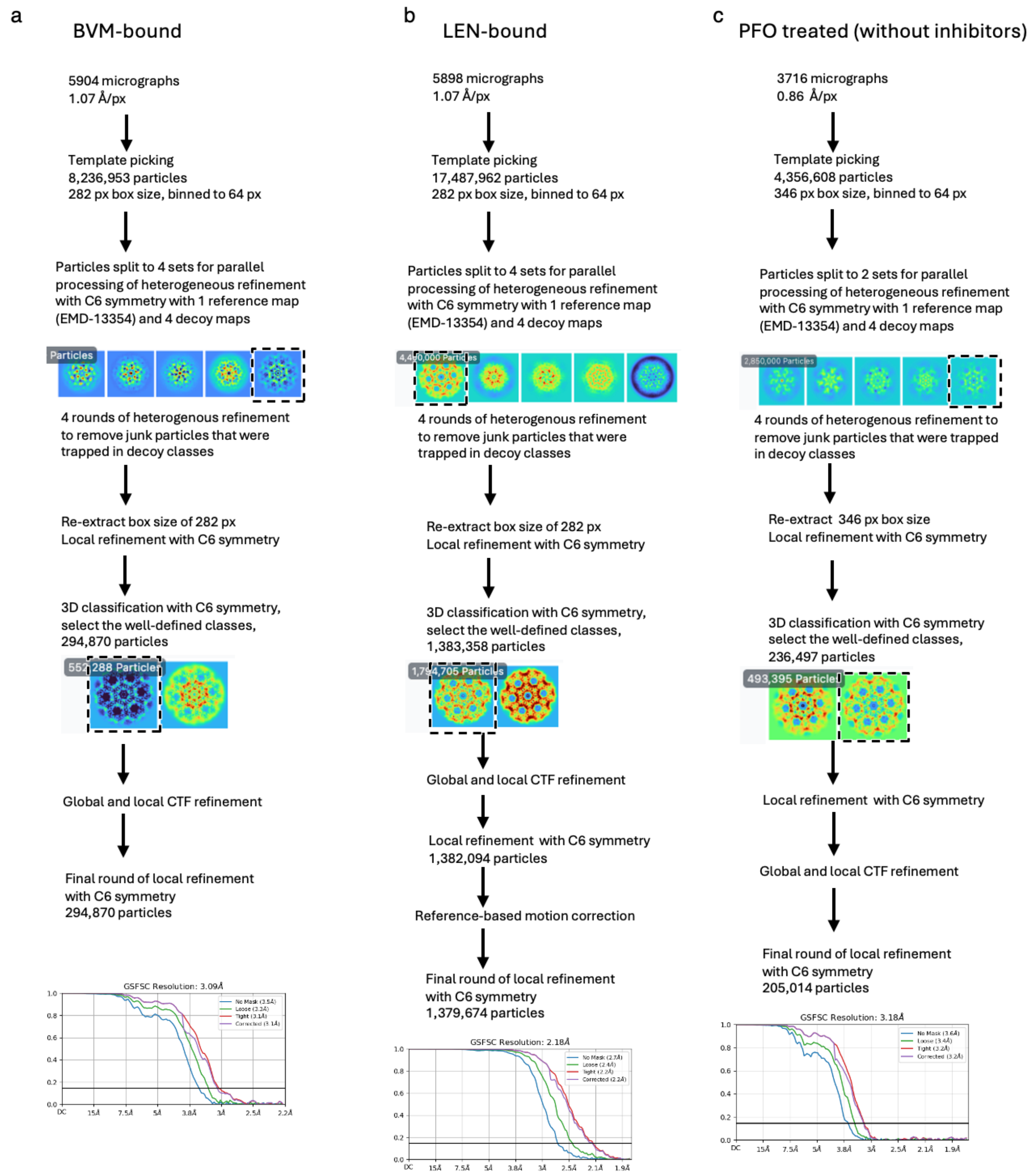


**Fig. S10 | Cryo-EM workflow for PFO- and/or ligand-treated VLPs**.

a. Data processing workflow for the reconstruction of the immature $Gag_{CA\text{-}SP1}$ with BVM-bound PFO-perforated NL-MA/NC VLPs.
b. Data processing workflow for the reconstruction of the immature $Gag_{CA\text{-}SP1}$ with LEN-bound PFO-perforated NL-MA/NC VLPs.
c. Data processing workflow for the reconstruction of the immature $Gag_{CA\text{-}SP1}$ from PFO-perforated NL-MA/NC VLPs.
